# Robust and Quality-of-Life-Aware Treatment Protocols in NSCLC using Deep Reinforcement Learning

**DOI:** 10.64898/2026.09.01.748242

**Authors:** Laura R. Jansén-Storbacka, Anne-Marie C. Dingemans, Kateřina Staňková, Alethea B. T. Barbaro, Sepinoud Azimi

**Affiliations:** Institute for Health Systems Science, Faculty of Technology, Policy and Management, Delft University of Technology, Delft, The Netherlands; Department of Respiratory Medicine, Erasmus Medical Center Cancer Institute, Rotterdam, The Netherlands; Delft Institute of Applied Mathematics, Delft University of Technology, Delft, The Netherlands

## Abstract

Under current systemic treatment of metastatic cancer, a drug is frequently prescribed at maximum tolerable dose (MTD) until either unacceptable toxicity or progression. Unfortunately, in many patients this treatment strategy leads to the development of treatment resistance. Evolutionary therapy approaches aim to forestall or delay treatment resistance in cancer by exploiting eco-evolutionary interactions. A well-known implementation is the adaptive therapy protocol of Zhang et al., in which tumour burden thresholds are used to guide strategic treatment holidays. Deep reinforcement learning (DRL) has recently been used to optimise these approaches. However, research combining DRL with evolutionary therapy approaches has so far focused on time to progression (TTP) as a performance metric, and has not quantified safety in terms of robustness to delayed treatment restart or included patient preferences regarding quality of life (QoL) in treatment design. In our study, we use a DRL agent informed by a mathematical two-population tumour growth model to design treatment schedules for patients with non-small cell lung cancer (NSCLC). The agent is trained on a virtual patient cohort using parameters previously fitted to data from patients with NSCLC treated with erlotinib. Beyond TTP, we focus on improving robustness to delayed treatment restart and on how individual preferences and values impact QoL experienced during treatment. We compare TTP, robustness and QoL under the DRL policy, the adaptive therapy protocol of Zhang et al., and MTD. We introduce a robustness metric “margin-to-failure” (MTF), and compare quality-adjusted-survival (QAS) across different patient preference profiles. Finally, we explore reward shaping to assess how QoL preferences can be incorporated into DRL-based treatment design. To evaluate our results, we consider different decision intervals, defined as the time between dosing adjustments. The DRL policy achieved greater median TTP, MTF, and QAS across all treatment decision intervals compared to the other two protocols. As decision intervals increased, TTP under DRL declined gradually towards that achieved under MTD. In contrast, the Zhang et al. protocol performed inconsistently and could result in premature progression. Additionally, a population-level policy trained on a cohort of virtual patients produced an interpretable treatment rule that extended TTP for most previously unseen patients and indicated that treatment should resume at a lower tumour burden when monitoring is less frequent. These findings show that DRL can balance the benefit of preserving drug-sensitive cells to suppress resistance against the risk of unsafe tumour regrowth. Reward shaping further showed how treatment strategies could be adjusted to reflect different patient preferences. Together, these results provide a biologically informed approach for designing robust and patient-centred evolutionary therapies in fast-growing cancers such as NSCLC.

## 1 Introduction

Treatment-induced resistance remains a major challenge in the care of advanced cancers [1, 9, 15]. This arises due to heterogeneity in the tumour microenvironment, where diverse cell populations compete for space and limited resources to proliferate and survive [20]. Under continuous maximum tolerated dose (MTD) therapy, treatment-sensitive cells are depleted, reducing competition for resources and allowing resistant mutants to proliferate (a phenomenon known as competitive release), ultimately leading to treatment failure [3, 10, 21, 25, 31]. Recognising cancer as an evolving Darwinian system has led to a new type of treatment strategy based on evolutionary game theory [27, 30, 31, 33, 40, 41, 43]. We use the term evolutionary therapy to refer to all strategies that explicitly account for and try to steer the ecological and evolutionary dynamics of tumour cell populations in order to delay or suppress the emergence of treatment resistance. Within this, we consider adaptive therapy to include strategies where treatment is decided (or adapted) in response to the current tumour state, and not according to a predetermined schedule such as in intermittent strategies. Evolutionary therapy approaches are often guided by mathematical models of the ecological and evolutionary dynamics. One clinically influential example in metastatic prostate cancer is the adaptive therapy protocol developed by Zhang et al. [42–44]. In this protocol, treatment is cycled on and off according to predefined tumour-burden thresholds, and tumour burden is monitored using PSA, a blood-based biomarker.

Prostate cancer is a relatively slow-growing cancer, and PSA measurements allow frequent monitoring of tumour dynamics. While the Zhang et al. protocol can maintain competition between treatment-sensitive and treatment-resistant populations under these conditions, longer monitoring intervals may allow substantial regrowth of the sensitive population between assessments, and the tumour burden to exceed the intended restart threshold [7]. In some patients, rapid tumour growth, combined with sparse measurements means that the Zhang et al. protocol may result in premature progression (progression that occurs earlier than would be expected under MTD). Non-small cell lung cancer (NSCLC) is the leading cause of cancer death [1], and represents an important example of such a setting [28]. NSCLC is a fast-growing, aggressive cancer which frequently presents with locally advanced or metastatic disease [14]. Because predictive serum-based biomarkers for NSCLC are scarce and unreliable [32], tumour burden in NSCLC is typically monitored using radiographic imaging performed every 6 *−* 12 weeks [28], with disease progression assessed according to RECIST criteria [4]. The combination of longer assessment intervals and fast growth creates challenges when applying the Zhang et al. protocol in NSCLC. Although this protocol can work under frequent monitoring [16], avoiding premature progression requires frequent monitoring of tumour burden [23, 29]. A subset of NSCLCs harbour actionable driver mutations, including EGFR mutations, that can be treated with targeted therapies such as erlotinib. These therapies are typically administered continuously at MTD. Under such treatment, sensitive cells are initially depleted, however, resistant cell populations subsequently expand and drive progression.

Although the Zhang et al. protocol may be challenging to implement given the current monitoring constraints of NSCLC, there is nevertheless a strong biological case for evolutionary therapies in this setting. This applies particularly in EGFR-mutated NSCLC treated with tyrosine kinase inhibitors (TKIs), where preclinical research suggests that resistant populations may grow more slowly than sensitive, and can even decline when treatment is paused, creating an opportunity to exploit competitive interactions between sensitive and resistant cancer cells through treatment holidays [2]. Although the underlying biology suggests evolutionary therapies may be beneficial in EGFR-mutated NSCLC, under sparse monitoring conditions the Zhang et al. protocol can be vulnerable to treatment delays. This clinical setting can be viewed as a sequential decision problem under incomplete state observation. Although the underlying tumour model contains treatment-sensitive and treatment-resistant populations, these subpopulations cannot be observed directly in clinical practice. Treatment decisions must instead be based on intermittent measurements of total tumour burden. This makes fixed threshold-based protocols vulnerable when tumour growth between assessments is rapid. In this study, we use virtual NSCLC patients to compare policies derived from deep reinforcement learning (DRL) against both continuous MTD therapy and the Zhang et al. protocol [10, 11, 37, 39].

Deep reinforcement learning (DRL) provides a way to learn treatment rules without prescribing fixed switching thresholds in advance. In DRL, an agent learns to maximise a reward by repeatedly interacting with an environment. At each decision time, the agent observes the current state, selects an action, and receives the reward. Recently, DRL has been applied to design evolutionary adaptive therapies in prostate cancer [6, 18, 19], non-cancer-specific chemotherapy [13], and, more recently, immunotherapy-treated NSCLC [17], while classical (tabular) RL methods have also been explored in breast cancer [34]. In our framework, the environment is provided by a two-population ordinary differential equation model of tumour growth, the observed state is the total tumour burden, and the available actions are treat at MTD or treatment holiday. The reward aims to encourage delayed progression while encouraging safe treatment holidays. The resulting DRL policy therefore depends on the specific tumour trajectories observed during training, rather than being constrained to predefined threshold rules. We train the agent on virtual patients with heterogeneous tumour growth and treatment-response dynamics, and evaluate the learned policies across clinically relevant decision intervals. We investigate whether DRL can learn dosing strategies that maintain a larger margin from progression and are less vulnerable to delayed treatment-restart decisions than the Zhang et al. protocol.

Existing reinforcement learning approaches to adaptive therapy primarily focus on tumour control metrics such as tumour burden or time to progression. This implicitly assumes that improved tumour control aligns with patient benefit, an assumption which may not hold in practice. Quality of life (QoL), which encompasses physical, psychological, and social well-being [35], is strongly influenced by treatment burden, i.e. the time and effort required for health care, and toxicity [24]. In EGFR-mutated NSCLC, prolonged TKI use is associated with cumulative toxicities [22, 26], highlighting the need to explicitly incorporate patient-centred outcomes into treatment design. QoL objectives are also individual, and patient preferences will determine the value of a treatment approach. Reinforcement learning is well suited to incorporating multiple objectives by including QoL directly within the reward function.

To address these challenges, we develop a DRL framework for EGFR-mutated, TKI-treated NSCLC that integrates tumour control with robustness and patient-centred objectives. Using a virtual patient cohort, we train a DRL agent to learn dosing policies under infrequent monitoring and compare its performance against an established adaptive protocol at both individual and population levels. We show how a population-trained policy increases time to progression for a majority of previously unseen virtual patients, suggesting a clinically feasible approach.

Finally, we extend the reward to incorporate explicit QoL preferences and examine how these reshape treatment strategies and trade-offs between survival, robustness, and patient well-being. Together, this provides a principled framework for designing adaptive therapies for NSCLC that are not only effective, but also robust and patient-centred.

## 2 Methods

### 2.1 Tumour growth model

Tumour dynamics were modelled using a two-population ordinary differential equation system describing interacting drug-sensitive and drug-resistant cells, with Gompertzian growth, asymmetric competition, and a log-kill treatment effect. The model structure and parameter distributions were derived from a previous analysis of START-TKI patients with stage III and IV erlotinib-treated NSCLC [16]. In that study, multiple models were fitted to longitudinal tumour burden data, and the model used here was identified as the best fitting among the candidate models.

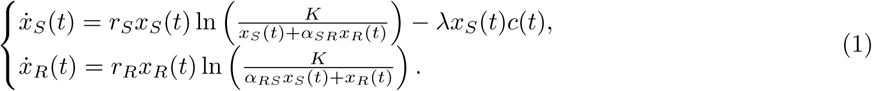

Here *x_S_* and *x_R_* denote sensitive and resistant populations, *r_S_,* and *r_R_* their growth rates, *K* the carrying capacity, *α_SR_,* and *α_RS_* the competition coefficients, *λ* the treatment effect, and *c*(*t*) the drug dose. The total tumour burden is *x*_total_(*t*) = *x_S_*(*t*) + *x_R_*(*t*). Dosing was modelled as a binary ON/OFF action. When dosing was on, medication was applied at constant MTD. This model captures the underlying eco-evolutionary tumour growth dynamics. The treatment effect reduces the sensitive population, while sensitive and resistant cell populations compete for space and resources through the shared carrying capacity. The competition coefficients allow this competition to be both asymmetric and frequency dependent, i.e. each cell type contributes differently to the effective population density which then affects the growth of the other type. While *α_SR_* weights the contribution of resistant cells to the competitive pressure experienced by sensitive cells, *α_RS_* weights the contribution of sensitive cells to the competitive pressure experienced by resistant cells.

### 2.2 Virtual patient cohort

To capture inter-patient heterogeneity, we generated a cohort of 60 virtual patients by sampling parameter sets for the tumour model (Equation 1), including growth rates, carrying capacity, drug effect, competition coefficients, and initial populations. Parameter distributions were obtained from the fitted parameter estimates reported in Jansén-Storbacka et al. [16]. Virtual patients were generated by sampling from these fitted parameter distributions. Parameters were sampled independently, with essential biological constraints enforced (e.g. *r_S_ > r_R_*), and weakly informative bounds used for interaction terms. Each sampled parameter set was screened for plausibility by simulating continuous treatment (MTD) and requiring an initial decline in tumour burden followed by regrowth after the nadir, producing the characteristic U-shaped trajectory associated with treatment-induced resistance. This filtering ensured that all virtual patients exhibited realistic treatment failure under MTD, avoiding ceiling effects in protocol comparison. Further details of parameter sampling, bounds, and filtering criteria are provided in the supplementary Section 1.

### 2.3 The clinical decision problem

Treatment decisions were made at discrete times *t_i_* = *i · d*, where *d* is the decision interval in days. Within each simulation and training run, *d* was fixed, so that *t_i_*_+1_ *− t_i_* = *d*. We evaluated policies separately for five decision intervals, *d ∈* {1, 14, 28, 42, 56} days. At each decision time, although the model contains both sensitive and resistant populations, the agent observes only the total tumour burden, in line with clinical reality where only imaging data is observed. Based on this observation, at time *t_i_* the agent selects either full-dose erlotinib (ON) or a drug holiday (OFF), and the action is held constant until the next decision at time *t_i_*_+1_. The progression endpoint and other evaluation metrics are defined in Section 2.5.

### 2.4 Benchmark treatment strategies

To evaluate the learned DRL policies, we compared them against standard clinical and theoretical treatment strategies. These benchmarks represent distinct philosophies of cancer control and provide interpretable reference points for evaluating learned policies.

#### Maximum tolerable dose (MTD)

MTD is the administration of the maximum tolerated dose until progression, and represents current clinical practice. In this work, constantly applied MTD is used as a baseline for comparison.

#### Zhang et al. protocol

We implemented the two-threshold adaptive therapy protocol of Zhang et al. as a clinically influential threshold-based benchmark [42, 43]. Let *x*_total_(*t*) = *x_S_*(*t*) + *x_R_*(*t*), and let *x*_total_(0) denote baseline tumour burden. Treatment was initially ON. At each decision time *t_i_*, the treatment action for the next interval was determined from the current tumour burden. Treatment was switched OFF if

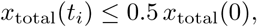

and switched ON if

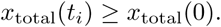

If tumour burden lay between these two thresholds, the previous treatment state was maintained. As for all intermittent policies in this study, ON corresponded to full-dose erlotinib and OFF corresponded to no treatment. The selected action was held fixed over the interval [*t_i_, t_i_*_+1_), and this cycle was repeated until either progression or the maximum simulation horizon was reached.

### 2.5 Evaluation metrics

All protocols were evaluated using time to progression (TTP), medication exposure (the proportion of time where medication is administered), and robustness to delayed treatment decisions.

#### Time to progression

TTP was defined as the first simulated time point at which the modelled total tumour burden, *x*_total_(*t*) = *x_S_*(*t*) + *x_R_*(*t*), reached or exceeded 20% above its baseline value:

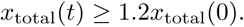

TTP was evaluated from the simulated tumour trajectory. This baseline relative approach to measuring progression is commonly used in evolutionary therapy modelling simulations where tumour regrowth is intentionally permitted during treatment holidays [6, 29, 42, 43].

#### Medication usage

Medication usage was defined as the percentage of simulated time during which treatment was ON. For each protocol, this was calculated as the total time receiving MTD divided by the total time until progression. Continuous MTD has medication exposure equal to 1, while the other protocols can have lower exposure depending on the duration and frequency of treatment holidays.

#### Premature progression

For protocol comparisons, we defined premature progression as progression occurring earlier than under continuous MTD treatment for the same virtual patient:

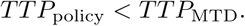

Differences of less than 1 day were treated as equivalent to MTD. In these cases, the evaluated policy reduced time to progression relative to the MTD baseline.

#### Margin-to-failure and Robustness to delayed treatment decisions

In addition to measuring time to progression (TTP), we quantified robustness to delayed treatment restart using a margin-to-failure (MTF) metric. For each virtual patient and protocol, we identified all scheduled treatment-restart decisions, defined as decision times at which treatment switched from OFF to ON. These OFF*→*ON switches are the points at which a delay would prolong a treatment holiday even when the protocol would have restarted medication.

At each scheduled restart point, we simulated a counterfactual delay scenario in which treatment remained OFF beyond the scheduled restart time. We then recorded the time from the scheduled restart decision until the total tumour burden reached the progression threshold, 1.2 *x*_total_(0). The patient-level MTF was defined as the minimum of these values across all OFF*→*ON switches for that patient and protocol. Intuitively, MTF represents the shortest delay in restarting treatment that could be tolerated before progression occurs. Larger MTF values indicate a wider safety margin and greater robustness to missed or delayed treatment restart. This metric is distinct from measurement error. Measurement error affects the observed tumour burden and can therefore cause incorrect or delayed switching decisions, whereas MTF quantifies how much delay in treatment restart could be tolerated once the protocol has indicated that treatment should be resumed.

To summarise robustness at the cohort level, MTF values were grouped into fragility categories:

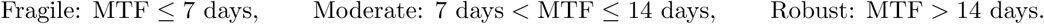

These thresholds correspond to one- and two-week delays, which reflect realistic time scales in calendar-based clinical care.

Two additional categories were used. If a protocol resulted in premature progression, then the case was labelled <u>Premature Progression</u>, because the protocol underperformed compared to the MTD baseline. Differences of less than one day relative to MTD were treated as MTD equivalent for classification purposes. If a policy used continuous MTD and therefore had no OFF*→*ON restart decisions, the case was labelled <u>Saturated</u>. MTF was computed for decision intervals *≥* 14 days. We did not compute MTF for *d* = 1 because daily decision making represents an idealised monitoring scenario rather than a clinically feasible imaging schedule in NSCLC. Additionally, the clinically interpretable fragility thresholds used here of 7 and 14 days represent 1 and 2 weeks delays, and for *d* = 1 only delays *≤* 1 day would be informative.

### 2.6 Reinforcement learning framework

Unlike the Zhang et al. protocol, which specifies fixed tumour-burden thresholds in advance, DRL learns a treatment policy directly from simulated tumour trajectories. In our framework, the agent observes tumour burden at each decision time, selects either full-dose erlotinib or a treatment holiday, and receives rewards designed to favour delayed progression and safe intermittent treatment. The following paragraphs define the environment, observed state, action space, reward function, learning algorithm, and training experiments.

Throughout this manuscript we distinguish between the learned <u>DRL policy</u> and the resulting <u>DRL protocol</u>. The DRL policy is the trained neural network that maps the observed tumour burden at a decision time to a probability of selecting treatment ON or OFF, while the DRL protocol is the treatment schedule generated when this policy is applied to a virtual patient. As each decision time the policy selects an action which is held fixed until the next decision time. The policy is therefore the learned decision rule of the neural network, while the protocol is its clinical application. For interpretability we summarise some policies by their treatment-propensity boundary [6], defined as the tumour burden when the probability of treatment equals 0.5, however, this boundary is an interpretable simplified summary of the policy, which is not equal to the full neural network.

#### 2.6.1 Environment and episodes

The DRL environment was defined using the coupled tumour-growth ODE system in Equation 1. At each clinical decision time *t_i_*, the agent selected an action, treatment ON or OFF, which was held constant over the interval [*t_i_, t_i_*_+1_). The ODE system was then integrated forward under this fixed action to generate the next observation. A training episode corresponds to one simulated treatment trajectory for a virtual patient, beginning at baseline and ending at progression or when the maximum analysis horizon is reached. Unless otherwise stated, each DRL agent was trained for 60,000 episodes.

##### Observed state

At each decision time *t_i_*, the agent observes the baseline-normalised total tumour burden, which is the clinically observable component of the tumour state:

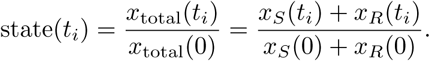

We restrict the state observed by the agent to the total tumour burden because CT imaging provides aggregate tumour burden but does not show sensitive and resistant subpopulations. Normalisation by the baseline tumour burden enables comparison across patients with different initial tumour sizes and makes policies more interpretable as they are represented as fractions of the baseline. This baseline-normalised quantity is not bounded by 1, values above 1 correspond to tumour burden above baseline, with progression defined at 1.2 times the baseline tumour burden.

##### Action space and transition dynamics

At each decision time *t_i_*, the agent selected one of two discrete actions:

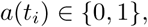

where *a*(*t_i_*) = 1 denotes full-dose erlotinib treatment (ON; 150 mg) and *a*(*t_i_*) = 0 denotes no treatment (OFF). The selected action was held fixed over the clinical decision interval [*t_i_, t_i_*_+1_), where *t_i_*_+1_ *− t_i_* = *d*. Between decision times, the coupled ODE system was integrated numerically using LSODA with adaptive time stepping and a maximum internal step size of 0.5 days.

#### 2.6.2 Reward Function

The reward function was designed to prioritise delayed progression while also encouraging treatment holidays only when the tumour burden remained under control. Following Gallagher et al. [6], the agent received a survival reward until progression and a terminal penalty at progression. Treatment holidays were rewarded if the tumour burden was below a safety threshold, while holidays above this threshold were penalised. In our implementation, rewards were scaled by the decision interval *d*, so reward magnitudes per-day were consistent across all decision intervals. We adapted the time-dependent long survival bonus to the shorter time-scale of NSCLC and applied it each year after 3 years. The default holiday threshold was set to baseline tumour burden, *x*_total_(0). Full numerical reward weights and hyperparameters are provided in Supplementary Section 2.

##### Learning algorithm

We trained the DRL agent using Advantage Actor-Critic (A2C). In this framework, the actor learns the treatment policy, i.e. the probability of selecting treatment ON or OFF from the observed tumour burden, while the critic learns to estimate the expected long-term reward of the current observed state. The actor and critic were implemented as two output heads of a shared neural network. Network architecture, loss function, optimiser settings, hyperparameter scaling, random seeds, and training and evaluation procedures are provided in Supplementary Section 2.

##### Starting point and NSCLC-specific adaptations

We followed the DRL architecture and reward setup of Gallagher et al. including (i) using only total tumour burden as the clinically observable input and (ii) using an LSTM-based actor-critic network to represent the treatment policy and value function. We modified this framework to account for the different growth and medication response dynamics of NSCLC treated with erlotinib [6, 8]. Key adaptations included scaling learning hyperparameters and reward magnitudes by the decision interval *d* to make training comparable across monitoring frequencies, and adjusting the timing of the long-term survival bonus start time to a clinically relevant timescale for NSCLC (survival beyond 3 years; see Subsection 2.6.2).

##### Training and evaluation experiments

Unless otherwise stated, separate DRL policies were trained for each virtual patient and decision interval.

To examine how monitoring frequency affects protocol performance in more detail, we performed an additional single patient sensitivity analysis. For one representative virtual patient, we trained and evaluated separate DRL policies for each integer valued decision interval from *d* = 1 to *d* = 84 days. The same patient was also simulated under MTD and the Zhang et al. protocol at each decision interval. This analysis was then used to examine how small changes in decision interval can alter performance under the different protocols, and to compare the interval sensitivity of the Zhang et al. and DRL protocols.

Following the cohort-training and fine-tuning strategy proposed by Gallagher et al. [6], we tested whether a population-trained DRL policy could provide a useful starting point for NSCLC treatment design. From the full cohort of 60 virtual patients, 30 were used for population-level training and the remaining 30 were held out for evaluation. At the start of each episode, one patient was sampled from the 30-patient training set, exposing the agent to heterogeneous tumour growth and treatment-response dynamics. The resulting population-level policy was evaluated without retraining on each of the 30 held-out patients. To assess whether this policy could also serve as a starting point for individualisation, we additionally fine-tuned it for each held-out patient by continuing training for 10,000 episodes from the population-trained parameters.

### 2.7 Quality of Life Components

The baseline reward prioritises survival and includes terms that discourage unnecessary treatment exposure, but it does not explicitly encode patient preferences. To explore how different patient priorities influence treatment recommendations, we define a QoL proxy comprising four clinically relevant burden components: tumour burden, treatment toxicity, medication burden, and monitoring burden, respectively. Tumour burden was defined as the total tumour burden relative to the progression threshold,

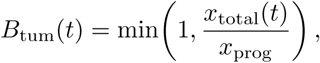

where *x*_prog_ = 1.2*x*_total_(0). Toxicity burden, *B*_tox_(*t*), was obtained from a one-compartment pharmacokinetic model with first-order elimination and a Hill-type mapping, scaled to lie in [0, 1]. Full details of the toxicity model and parameter defaults are provided in Supplementary Section 3. Medication burden was defined from the treatment action,

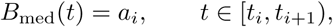

where *a_i_*= 1 denotes treatment ON and *a_i_*= 0 denotes treatment OFF. Monitoring/visit burden was defined as

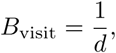

where *d* is the decision interval in days. More intensive monitoring therefore corresponds to a higher visit burden. Because *d* is fixed in this setup, *B*_visit_ is constant for each simulation.

#### Conversion to positive QoL components

To make interpretation more intuitive with higher values representing greater QoL, each burden was converted into a positive QoL component,

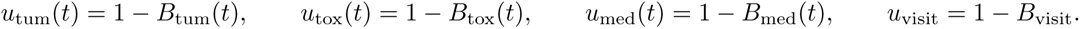

These components are bounded between 0 and 1, with larger values representing better quality of life. For example, *u*_tum_(*t*) decreases as tumour burden approaches the progression threshold, *u*_med_(*t*) = 1 during treatment holidays, and *u*_visit_ is higher when monitoring is less frequent.

#### Toxicity and medication represent distinct treatment-burden dimensions

Although toxicity and medication are correlated because toxicity is driven by drug exposure in the pharmacokinetic model, they represent distinct constructs. Medication burden captures the burden of being on treatment, including daily pill-taking, treatment routines, and the perceived burden of continuous medication, which may persist even when side effects are mild or absent. In contrast, toxicity represents physiological side effects from drug exposure, and is calculated using the pharmacokinetic toxicity model described in Supplementary Section 3.

##### 2.7.1 Preference-weighted QoL score

The four QoL components were combined using a patient-specific weight vector

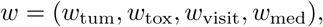

satisfying

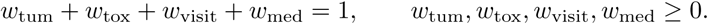

The resulting model-based QoL score was defined as

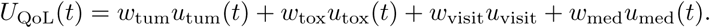

##### 2.7.2 Quality-adjusted survival (QAS)

Quality-adjusted survival (QAS) combines survival time with a time-varying quality-of-life weight [38]. In our framework, we define QAS as the time integral of the preference-weighted QoL score,

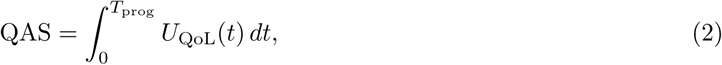

where *T*_prog_ denotes time to progression. In practice, the QAS integral was evaluated on the fine toxicity simulation grid. In our simulations, we approximated the integral using the trapezoidal rule with a default step size of (0.1) days. Treatment action (and thus medication utility) was piecewise constant within each decision interval [*t_i_, t_i_*_+1_), while visit utility remained constant within each simulation because the decision interval (d) was fixed. Toxicity was simulated directly on the fine grid, and tumour burden was mapped to that grid from the simulated trajectory (via interpolation).

QAS is conceptually related to quality-adjusted life-year calculations, but here QAS is reported in days and uses the model-based QoL proxy defined above rather than a validated health-utility instrument. We did not apply discounting, because the objective was to compare simulated treatment protocols over the same horizon rather than to perform a health-economic valuation.

We also report mean QoL score, QAS*/T*_prog_, to separate average quality of life during survival from survival duration. Protocols are compared at the paired patient level using differences in QAS, summarised by median and interquartile range, with significance assessed using two-sided sign tests. Details are provided in the Supplementary Section 7.

###### Preference profiles

To illustrate how patient priorities affect QoL-adjusted outcomes, we define five example preference profiles: balanced, medication-adverse, toxicity-adverse, tumour-burden-adverse, and visit-adverse (Table 1). We use these profiles to evaluate the existing protocols according to their QAS performance.

**Table 1:**
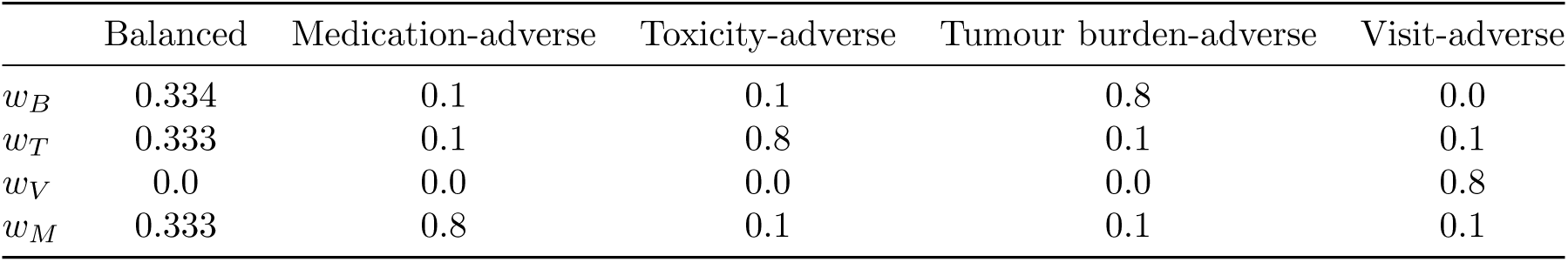
Preference profiles used for QoL evaluation.

|  | Balanced | Medication-adverse | Toxicity-adverse | Tumour burden-adverse | Visit-adverse |
| --- | --- | --- | --- | --- | --- |
| $w_B$ | 0.334 | 0.1 | 0.1 | 0.8 | 0.0 |
| $w_T$ | 0.333 | 0.1 | 0.8 | 0.1 | 0.1 |
| $w_V$ | 0.0 | 0.0 | 0.0 | 0.0 | 0.8 |
| $w_M$ | 0.333 | 0.8 | 0.1 | 0.1 | 0.1 |

Medication burden and toxicity were kept as separate preference dimensions because they represent distinct aspects of the patient experience. Medication burden captures the burden of being on treatment, whereas toxicity captures physiological side effects from drug exposure. In the present binary ON/OFF dosing framework these quantities are closely correlated, because toxicity is driven primarily by cumulative drug exposure. However, they need not coincide in more general settings, for example if multiple dose levels, patient-specific toxicity sensitivity, or alternative drugs are considered.

Visit burden was included to evaluate how preferences about monitoring frequency affect QAS across different decision intervals. In this study, the monitoring interval *d* was fixed before each simulation and was not chosen by the DRL policy. Thus, the treatment policy determines whether medication is ON or OFF, but it does not determine visit frequency. The visit-adverse profile therefore evaluates the value of less frequent monitoring as an externally specified clinical preference rather than as an optimised policy decision. In the other profiles, the visit-burden weight was set to zero so that those profiles focus on tumour burden, toxicity, and medication burden.

##### 2.7.3 Exploring Quality-of-life aware reward shaping

In addition to evaluating the baseline DRL, Zhang et al.protocol, and MTD using QAS, we explored whether reward shaping (modifying the reward function weights and thresholds) could push the learned policies towards prioritising different patient QoL preferences. For example, a lower holiday threshold could encourage earlier treatment resumption in a tumour-burden-adverse setting.

###### Reward-parameter sweep

We performed an exploratory reward-parameter sweep for decision intervals *d ∈ {*14, 28, 42*}* days in two representative virtual patients. For each reward configuration, a separate DRL agent was trained and evaluated. The swept parameters included the holiday threshold, holiday penalty, holiday bonus, and survival reward. The full set of parameter ranges and training details used in the sweep are provided in Supplementary Section 2.6. The sweep was used to identify how different reward components affected learned treatment behaviour. Lower holiday thresholds and stronger holiday penalties tended to promote earlier treatment resumption, corresponding to a more tumour-burden-adverse strategy. Higher holiday thresholds and larger holiday bonuses tended to encourage longer or more frequent treatment holidays, corresponding to more medication- or toxicity-adverse strategies. Increasing survival-related rewards tended to prioritise longer tumour control. These mappings were used to interpret the resulting survival-QoL trade-offs, rather than as a set of pre-specified preference-specific reward functions.

## 3 Results

### 3.1 DRL protocols maintain higher median TTP across decision intervals

Across all evaluated decision intervals, the DRL protocol achieved higher median TTP compared to both the MTD and Zhang et al. protocols. Under continuous MTD, TTP is 15.11 months for all decision intervals. At shorter decision intervals both the DRL and Zhang et al. protocol improved TTP compared to MTD. When *d* = 1, the median TTP was 49.59 months under the DRL, and 40.41 months under the Zhang et al. protocol, while at *d* = 14, median TTP under the DRL policy was 35.66 months compared to 29.46 for the Zhang et al. protocol.

For longer decision intervals of *d* = 28*−*56, the median DRL performance declined gradually but remained above MTD with median TTP values of 25.85, 21.42, and 18.95 months at *d* = 28, 42 and 56 respectively. In contrast median TTP under the Zhang et al. protocol decreased below MTD at longer intervals, with values of 7.31, 5.75 and 5.79 months for *d* = 28, 42 and 56 respectively, indicating it is more sensitive to sparse monitoring. Premature progression under the Zhang et al. protocol often happens with longer decision intervals after only 1 or 2 ON-OFF medication cycles, due to fast growth of the sensitive population which then passes the progression threshold before a scheduled measurement and treatment decision. Detailed summary statistics are shown in the Supplementary Table S3.

Median TTP was significantly higher (*p ≤* 0.0001 for DRL compared to MTD across all decision intervals. The pairwise differences between the Zhang et al. protocol and MTD are only significant for *d* = 1 and *d* = 14, at longer intervals *d ∈* 28, 42, 56 they do not show a significant difference. Similar variability occurs in pairwise comparisons of the DRL and Zhang et al. protocols, with significant differences only for *d* = 1 and *d* = 56. This occurs due to inter-patient variability, where the Zhang et al. protocol gives longer TTP for some patients compared to MTD, but performs worse for others.

Kaplan–Meier analyses supported these trends (Figure 2). Pairwise log-rank tests showed significantly longer TTP for DRL compared to MTD for all decision intervals. Differences between the Zhang et al. protocol and MTD were only significant at *d* = 1 and *d* = 14 days, while comparisons between DRL and the Zhang et al. protocol were significant at *d* = 1 and *d* = 56 days. Kaplan–Meier curves for all decision intervals, together with their associated risk tables, are provided in Supplementary Figure S2, and pairwise log-rank test results are reported in Supplementary Table S4.

**Figure 1:**
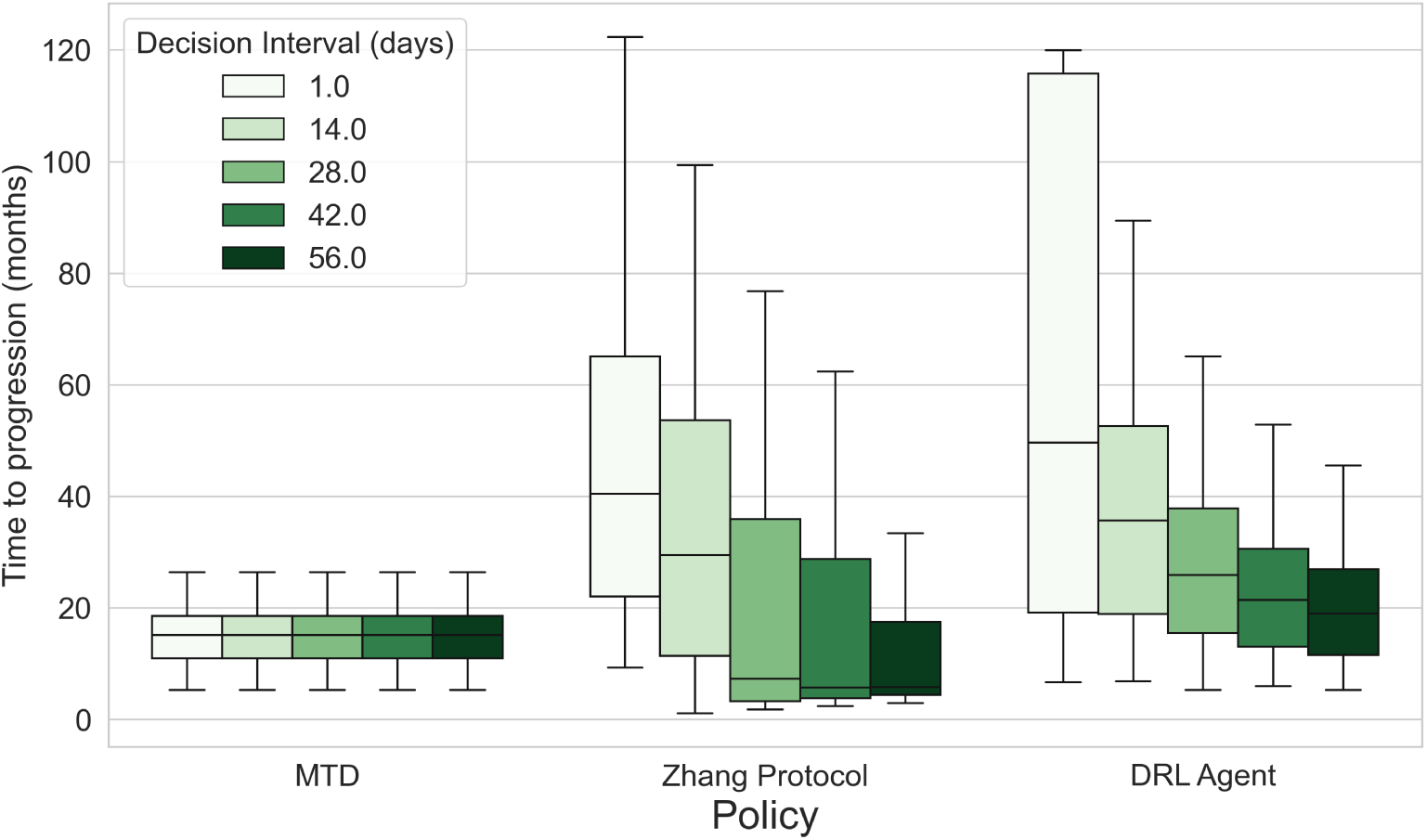
DRL protocols maintain higher TTP than MTD under sparse decision making. Box plots showing time to progression (TTP) for MTD, the Zhang et al. protocol, and the DRL policy across decision intervals of 1, 14, 28, 42, and 56 days. Horizontal lines indicate medians, boxes show the interquartile range (IQR), and whiskers extend to 1.5*×* the IQR. MTD gives the same TTP across decision intervals because treatment is continuous. The Zhang et al. protocol improves median TTP relative to MTD at short decision intervals (1 or 14 days), but gives lower TTP than MTD at longer intervals (28,42 and 56 days) because sparse monitoring can lead to premature progression. In contrast, DRL maintains median TTP above MTD across all evaluated decision intervals, although performance declines towards MTD as decisions become less frequent. Detailed summary statistics are provided in Supplementary Table S3.

**Table 2:** Premature progression (PP) relative to standard-of-care continuous MTD. Counts indicate the number of virtual patients (out of *N* = 60 per decision interval) with premature progression, defined as TTP_policy_ *<* TTP_MTD_ day. For these cases, underperformance is reported as the median and range of the TTP loss relative to MTD (days).

| Decision (days) | Zhang et al. |  | DRL |  |
| --- | --- | --- | --- | --- |
| | PP $n$ (%) | TTP loss | PP $n$ (%) | TTP loss |
| 1 | 0 (0.0%) | – | 0 (0.0%) | – |
| 14 | 15 (25.0%) | 201.3 [23.7–497.0] | 0 (0.0%) | – |
| 28 | 33 (55.0%) | 237.9 [59.1–525.6] | 0 (0.0%) | – |
| 42 | 35 (58.3%) | 209.0 [3.7–578.8] | 0 (0.0%) | – |
| 56 | 41 (68.3%) | 239.7 [9.7–645.9] | 1 (1.7%) | 14.1 [14.1–14.1] |

### 3.2 The Zhang et al. protocol is highly sensitive to decision interval under sparse monitoring

Although the Zhang et al. protocol improved median TTP compared to MTD under frequent monitoring, its performance became highly sensitive to decision interval with more cases of premature progression as monitoring became less frequent. This is consistent with previous work showing that the Zhang et al. protocol can improve tumour control when treatment decisions are made sufficiently frequently [16, 37]. The crossing Kaplan–Meier curves (Figure 2) for the Zhang et al. and MTD protocols at decision intervals *d ∈* 14, 28, 42, 56 suggest variable patient-level effects, with some virtual patients benefiting from treatment holidays and others progressing earlier than under MTD. As the decision intervals increase, the crossing points between the Zhang et al. protocol occurs at progressively lower survival probabilities, indicating that an increasing larger fraction of patients under the Zhang et al. protocol show premature progression i.e. shorter TTP compared to the MTD protocol.

**Figure 2:**
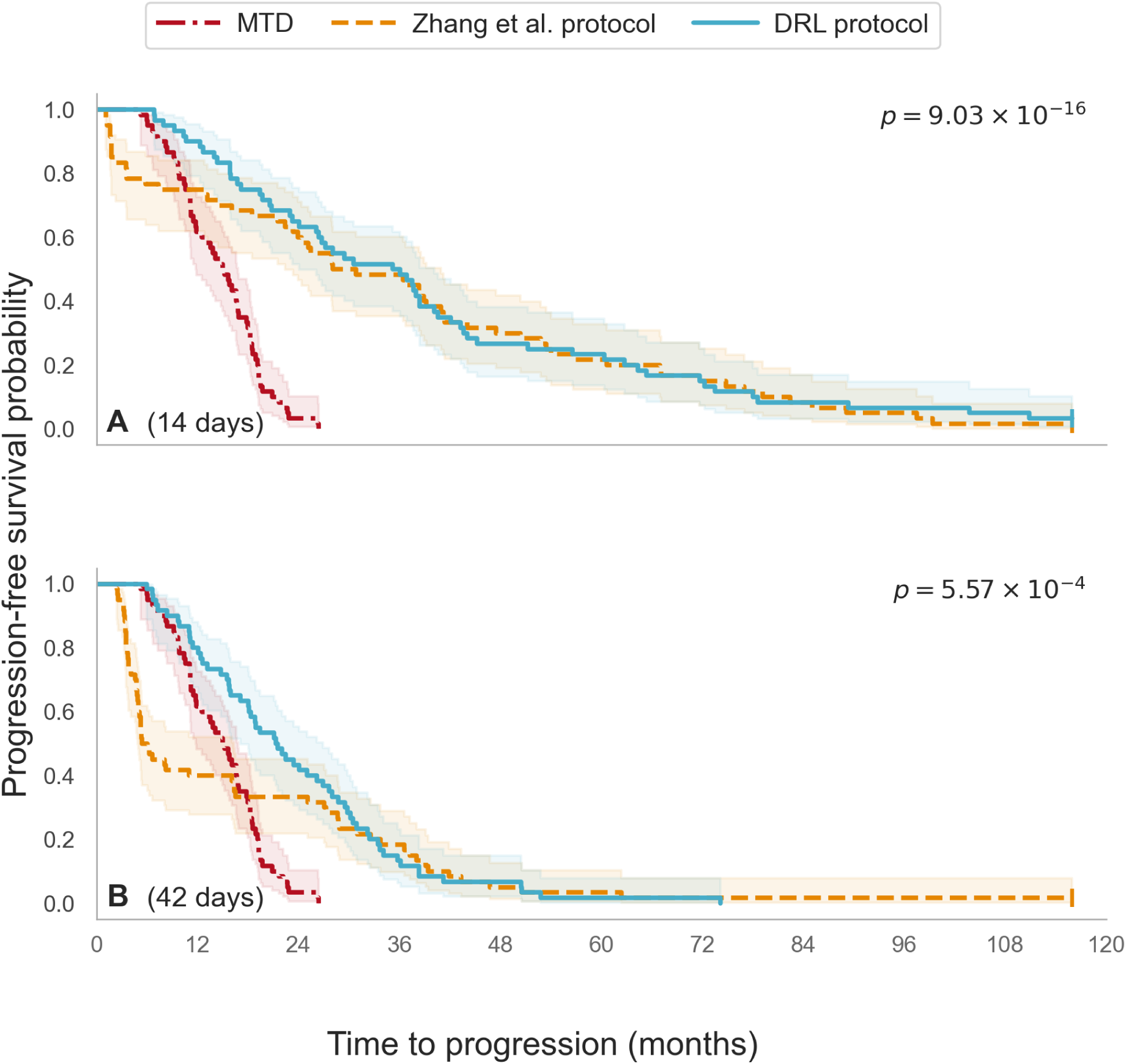
Kaplan–Meier estimates of time to progression (TTP) by treatment protocol for 14-day and 42-day decision intervals. **A,** 14 days; **B,** 42 days. Curves are shown for MTD, Zhang et al., and DRL protocols. Global *p*-values from log-rank tests indicate at least one significant difference. Corresponding pairwise *p*-values are reported in Supplementary Table S4. and Kaplan-Meier curves for all decision intervals plus risk tables are shown in the Supplement in Figure S2. Counts of cases where a policy predicts lower TTP than MTD, grouped by decision interval are summarised in Table 2.

Across all patient–interval evaluations, the Zhang et al. protocol resulted in premature progression in 41.3% of virtual patients, with the median reduction in TTP exceeding 200 days for *d ≥* 14. In contrast, the DRL protocol showed premature progression in only 1/300 patient–interval evaluations, with a TTP reduction of 14.1 days at *d* = 56. Inspection of the trajectory showed that the DRL protocol discontinued treatment after competitive release had already occurred, accepting a small reduction in TTP in exchange for additional treatment-holiday reward. Full premature progression counts and ranges for all decision intervals are shown in Table 2.

In contrast to the gradual decline in TTP observed under DRL as decision interval increased, the Zhang et al. protocol performance varies, with transitions between tumour control and premature progression as the decision interval changes. This interval-sensitivity is consistent with Zhang et al., who showed that adaptive-treatment cycle lengths depend on patient-specific tumour composition and competitive interactions between subpopulations [43].

The Zhang et al. protocol checks treatment-restart thresholds only at discrete decision times. During treatment holidays, the sensitive population can regrow; while this may help maintain competition against resistant cells, it also increases total tumour burden. If the decision interval is too long for a given patient’s growth dynamics, tumour burden can cross the intended restart threshold between assessments while treatment remains OFF. In patients with fast-growing cancers, the tumour burden can cross progression threshold before the treatment is resumed.

In the more aggressive NSCLC there are many fast-growing patients and small changes in monitoring frequency can mean the difference between tumour control and premature progression. This instability arises from the combination of decision intervals and individual growth dynamics, meaning an interval that can control tumour burden in one patient may lead to premature progression in another. This interval-dependent behaviour is demonstrated in detail for a single patient in Figure 3.

**Figure 3:**
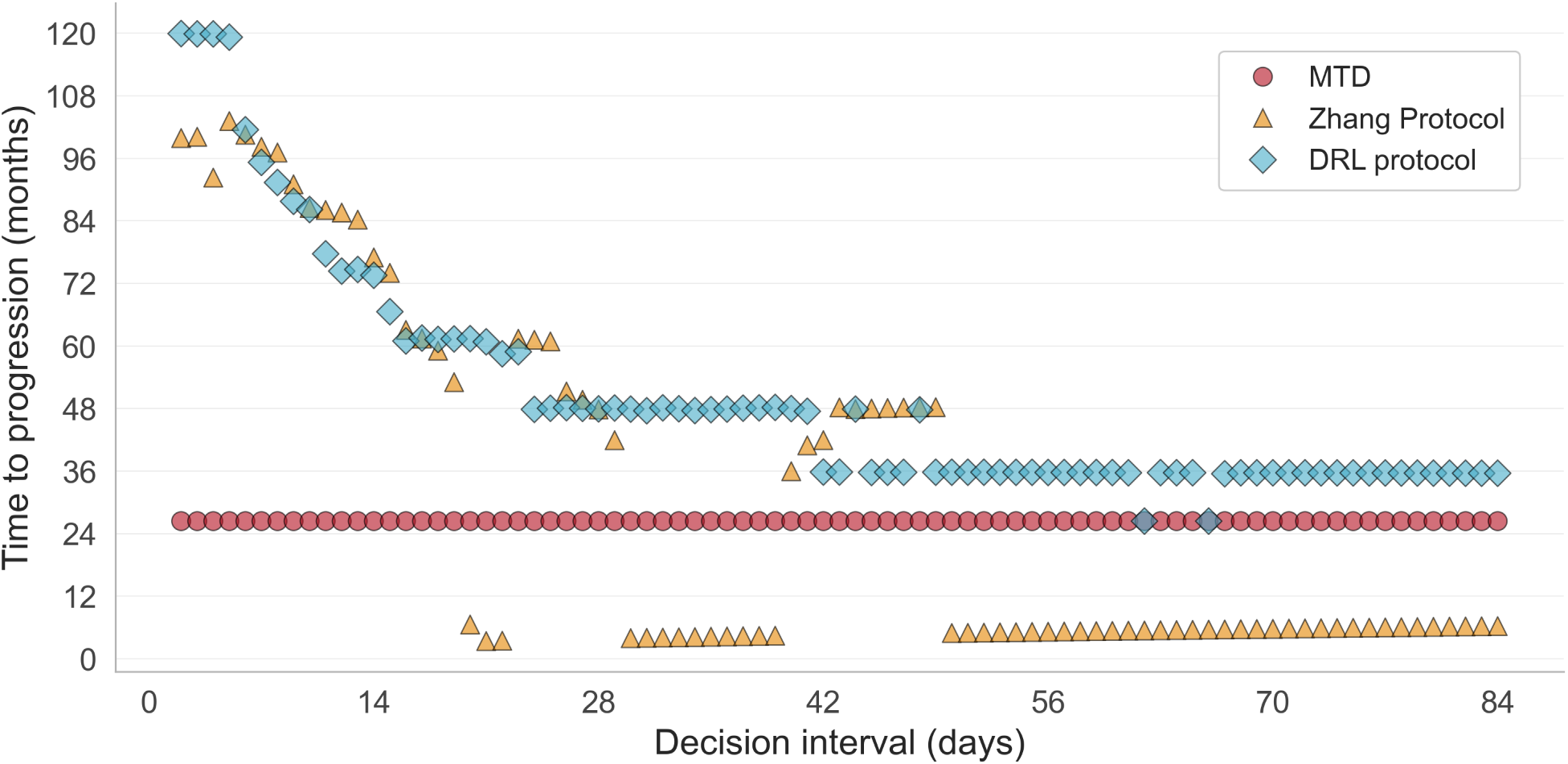
Single-patient sensitivity to decision-interval length. Time to progression (TTP) for one representative virtual patient under MTD, the Zhang et al. protocol, and the DRL policy as a function of decision-interval length, from 1 to 84 days. Progression was defined as the first simulated time point at which total tumour burden exceeded 1.2 *x*_total_(0). This high-resolution interval sweep was performed for one patient to illustrate the interval sensitivity observed at the cohort level (see Table 2). The Zhang et al. protocol shows abrupt changes in TTP across nearby decision intervals, alternating increased TTP relative to MTD and premature progression. In contrast, the DRL policy changes more gradually as the decision interval increases, and still learns policies that include safe treatment holidays.

Together, these results show that the Zhang et al. protocol is highly sensitive to the timing of threshold checks. Because treatment restart is evaluated only at discrete decision times, small changes in the decision interval can determine whether treatment restart happens before progression occurs. In fast-growing tumours such as NSCLC, a small change can shift the outcome from control to premature progression. In contrast, the learned DRL policy changes more gradually as monitoring frequency decreases, shifting towards MTD rather than learning policies that cause premature progression.

### 3.3 DRL policies are more robust to delayed treatment restart

We next used MTF to evaluate robustness to delayed treatment restart, with lower MTF indicating a narrower safety margin. Although DRL achieved higher median TTP than the Zhang et al. protocol at the population level, this was not true for every individual patient. We therefore examined whether cases with longer TTP under the Zhang et al. protocol also maintained a clinically useful safety margin to delayed treatment restart. This is illustrated for one virtual patient in Figure 4. For this patient, the Zhang et al. protocol achieved longer TTP than the DRL policy (2343 vs 2238.5 days), but had a much smaller safety margin (MTF of 1.3 vs 10.4 days). This means the Zhang et al. protocol achieved the longer simulated survival by restarting treatment very close to progression, whereas the DRL policy sacrificed a small amount of TTP to maintain a larger buffer against delayed restart.

**Figure 4:**
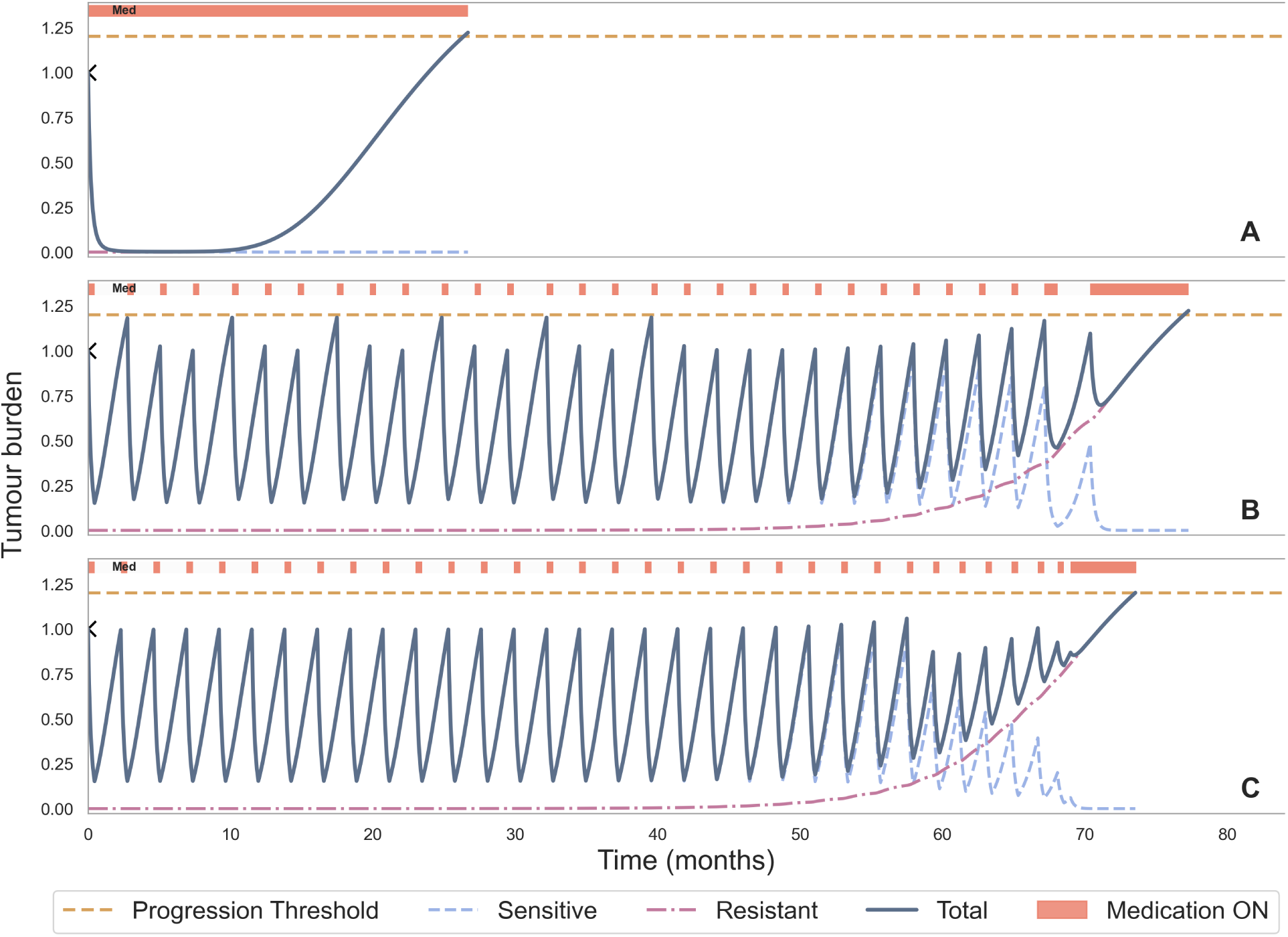
Example of robustness trade-off with *d* = 14 days. Simulated trajectories under **(A)** MTD, **(B)** the Zhang et al. protocol and **(C)** the DRL learned policy for one representative virtual patient. Although the Zhang et al. protocol achieves longer TTP than the DRL in this example, it does so with a much smaller margin-to-failure (MTF 1.276 vs 10.359 days). MTF, defined as the shortest time from any scheduled OFF*→*ON restart decision to progression under a counterfactual scenario in which treatment remains OFF, was computed at scheduled OFF*→*ON treatment-restart decisions. For the Zhang et al. protocol, delaying the most vulnerable scheduled restart by 2 days would result in progression, whereas the DRL policy maintains a larger safety margin of more than 7 days while still increasing TTP relative to MTD.

At the population level, the Zhang et al. protocol frequently produced cases with narrow safety margins, with many cases having MTF *<* 7 days (Figure 5). In contrast, DRL policies more often maintained larger MTF values, indicating a wider buffer before progression if treatment restart were delayed. This difference is also visible across decision intervals (Figure 6), as the DRL has fewer fragile and premature progression cases compared to the Zhang et al. protocol.

**Figure 5:**
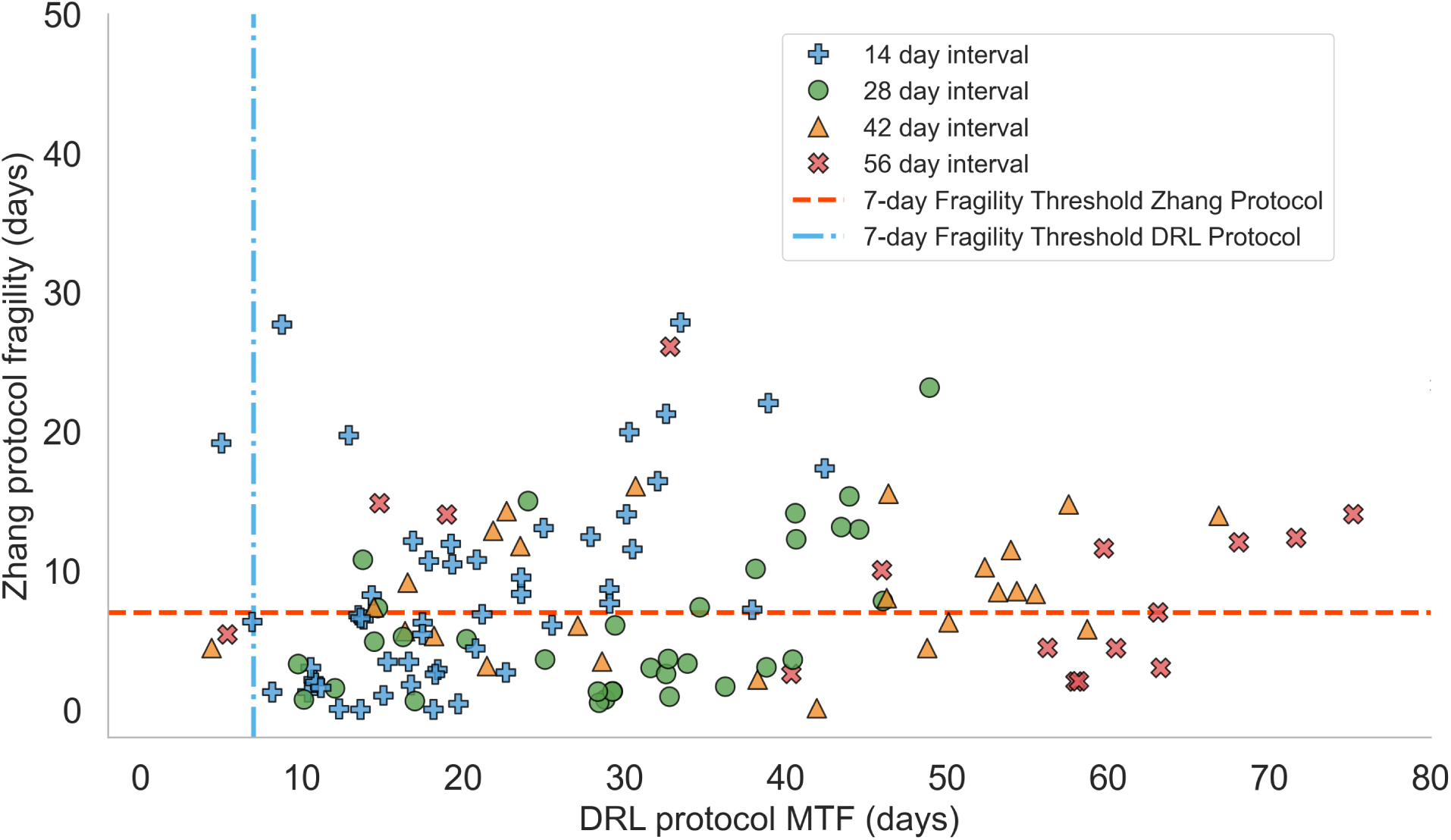
Patient-level robustness comparison (MTF) between the Zhang et al. protocol and DRL. Scatterplot of margin-to-failure (MTF) values for individual virtual patients comparing the Zhang et al. protocol versus DRL at matched decision intervals. Each point corresponds to one patient interval evaluation with defined MTF for both protocols (cases with undefined MTF are excluded; see Figure 6 for “Premature progression” and “Saturated” categories). Points below the horizontal line indicate a MTF of *<* 7 days for the Zhang et al. protocol, while points to the left of the vertical line indicate the same for the DRL protocol.

**Figure 6:**
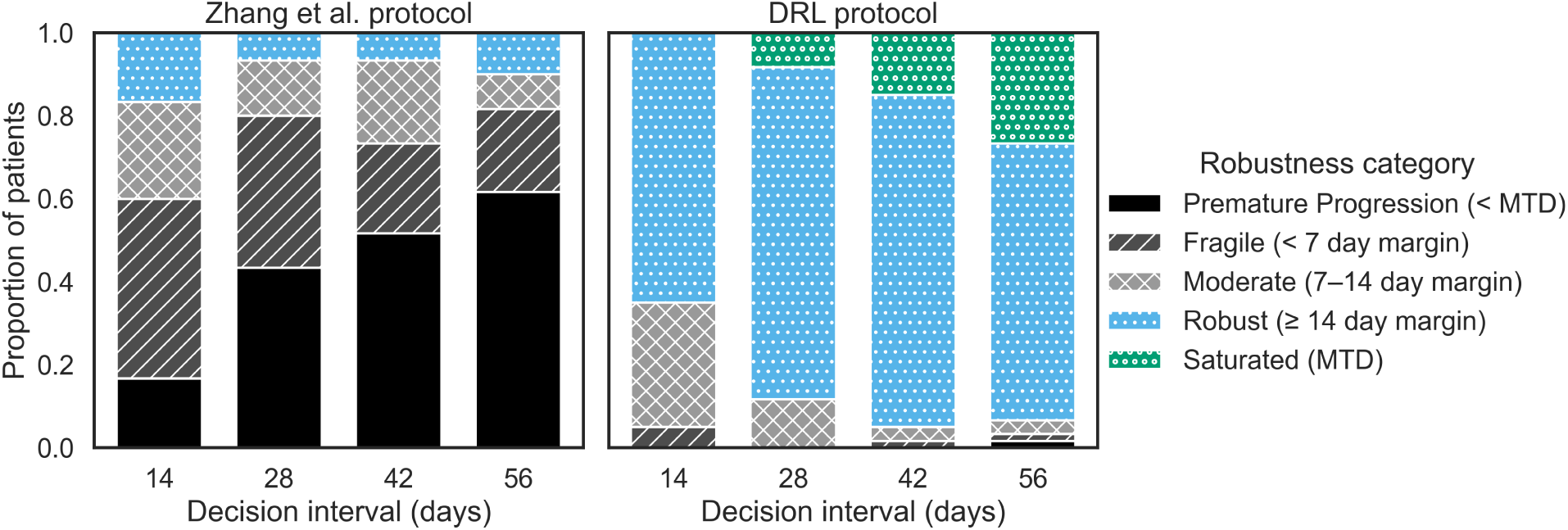
Robustness of DRL and Zhang et al. protocols across decision intervals. Stacked bar plots show the proportion of patients in each MTF robustness category for each protocol and decision interval. Fragile, Moderate, and Robust correspond to MTF values of *≤* 7, *>* 7 to *≤* 14, and *>* 14 days, respectively. Premature progression indicates cases where the protocol reduced TTP relative to MTD by more than 1 day. Saturated indicates cases that used continuous treatment or produced an outcome effectively equivalent to continuous treatment, and therefore had no informative OFF*→*ON restart decisions. Premature progression and Saturated cases are not assigned MTF values. DRL shows fewer fragile and premature-progression cases than the Zhang et al. protocol across decision intervals, with saturated cases becoming more frequent at longer decision intervals.

At longer decision intervals, some DRL policies become saturated and the agent chooses MTD. Biologically, this can occur when the treatment effect is strong enough so that the initial treatment ON period depletes the sensitive population before a holiday is possible. This removes the competitive control of resistant cells that treatment holidays are designed to preserve. After this happens TTP is decided by the resistant growth rate alone and is equivalent to MTD. Alternatively, in cases with a fast sensitive growth rate, the sparse monitoring may allow for substantial regrowth and lead to premature progression during treatment holidays. In this case, the DRL agent learns to choose MTD to maximise TTP as treatment holidays always lead to premature progression. Consequently, the improved robustness of the DRL compared to the Zhang et al. protocol comes partly at the cost of higher medication exposure as decision intervals become longer.

Improvements in robustness at longer decision intervals are accompanied by increased medication exposure (Figure 7). At longer intervals, the DRL policy more frequently approaches continuous treatment in a subset of patients, as reflected by the increase in saturated cases. This highlights a trade-off between maintaining tumour control and minimising treatment burden, which is captured in the QoL framework.

**Figure 7:**
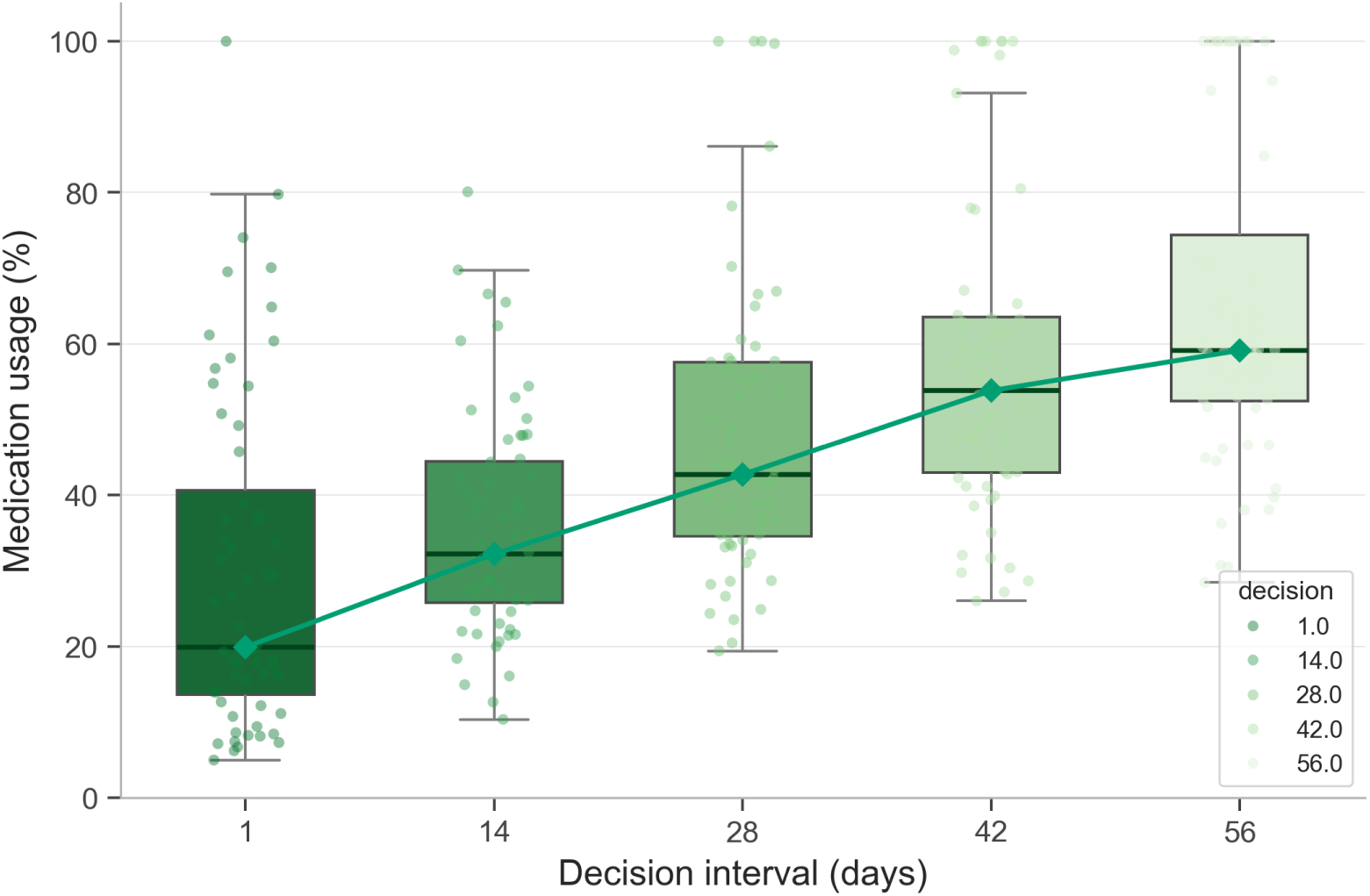
Medication use increases with decision interval under the DRL policy. Boxplots show the population distributions of the percentage of time that medication is ON at each decision interval. Boxes indicate the interquartile range, horizontal lines within boxes indicate the median, whiskers extend to the most extreme values within 1.5 times the interquartile range, gray points indicate individual patients, and the connected green diamonds indicate the median trend across decision intervals.

### 3.4 Population-trained policies generalise to unseen patients and suggest feasible treatment boundaries for NSCLC

Previous work in prostate cancer showed that DRL policies with binary ON/OFF action spaces can often be summarised using treatment-propensity curves, which relate tumour burden to the probability of choosing ON [6]. In this interpretation, the treatment-propensity boundary, above which medication should be turned on is the tumour burden at which *p*_treat_ = 0.5. Here, we extend this by quantifying the distributions of thresholds across patients and decision intervals in NSCLC.

Across individually trained policies, the treatment-propensity boundary exhibits substantial inter-patient variability at every decision interval. Nevertheless, there is a clear interval-dependent trend as the median treatment propensity boundary decreases with sparser monitoring, declining from 0.939 at *d* = 1 to 0.387 at *d* = 56 (Table 3). The distributions of the inferred boundaries also shifted towards lower tumour-burden values as the decision intervals increased with ranges shifting from 0.231–1.166 at *d* = 1 to 0.051–0.968 at *d* = 56, while the interquartile range changed from 0.534–1.046 to 0.276–0.528. This indicates that although individual treatment thresholds are patient dependent, the DRL policies consistently identify lower restart boundaries as monitoring becomes sparser.

**Table 3:** Treatment-propensity boundary summary statistics by decision interval. The boundary is defined as the baseline-normalised tumour burden at which the learned policy selects treatment ON with probability *p*_treat_ = 0.5..

| Decision (days) | Mean | Std | Median | Q25 | Q75 | Min | Max |
| --- | --- | --- | --- | --- | --- | --- | --- |
| 1 | 0.812 | 0.304 | 0.939 | 0.534 | 1.046 | 0.231 | 1.166 |
| 14 | 0.726 | 0.204 | 0.768 | 0.620 | 0.857 | 0.173 | 1.095 |
| 28 | 0.578 | 0.239 | 0.606 | 0.379 | 0.773 | 0.085 | 1.162 |
| 42 | 0.489 | 0.247 | 0.484 | 0.291 | 0.673 | 0.091 | 1.154 |
| 56 | 0.410 | 0.202 | 0.387 | 0.276 | 0.528 | 0.051 | 0.968 |

Although patient-specific DRL policies provide personalised treatment rules within our framework, they may not be feasible to train or update in practice. Such policies require patient-specific tumour dynamics and sufficient longitudinal observations, which present challenges in NSCLC where imaging is sparse and tumour growth can be rapid [28]. Previous work has proposed using pre-trained population based DRL models as an initial treatment policy which is later personalised as more patient specific information becomes available [6]. However in NSCLC the combination of fast dynamics and sparser observations mean there may not be time to calibrate individual dynamics as progression could occur before a single ON-OFF treatment cycle. We therefore asked whether population-level DRL policies could themselves provide clinically useful containment rules.

We trained a single population-level DRL policy and evaluated it on 30 unseen virtual patients. As with the patient-specific DRL policies, the population-trained policy could also be summarised by a treatment-propensity boundary, defined as the tumour burden at which *p*_treat_ = 0.5. As with the individually trained policies, the population-level threshold also declines as the decision interval increases, as illustrated for *d* = 14 and *d* = 28 in Figure 8. Because this policy was trained to perform across a heterogeneous cohort rather than individually for each held-out patient, it was not expected to improve TTP for every individual. When applied directly, without retraining, the population-trained policy improves time to progression relative to MTD for the majority of patients, with twenty-nine out of thirty (96.7%) patients at *d* = 14 and twenty-seven out of thirty (90%) patients at *d* = 28 achieving TTP_DRL_ *>* TTP_MTD_. Although performance was slightly reduced compared with individually trained policies, these results show that a single population-trained decision boundary can retain clinical benefit for most patients without patient-specific retraining. This contrasts with the Zhang et al. protocol at comparable intervals with 25% showing premature progression at *d* = 14 and 55% at *d* = 28 (Table 2).

**Figure 8:**
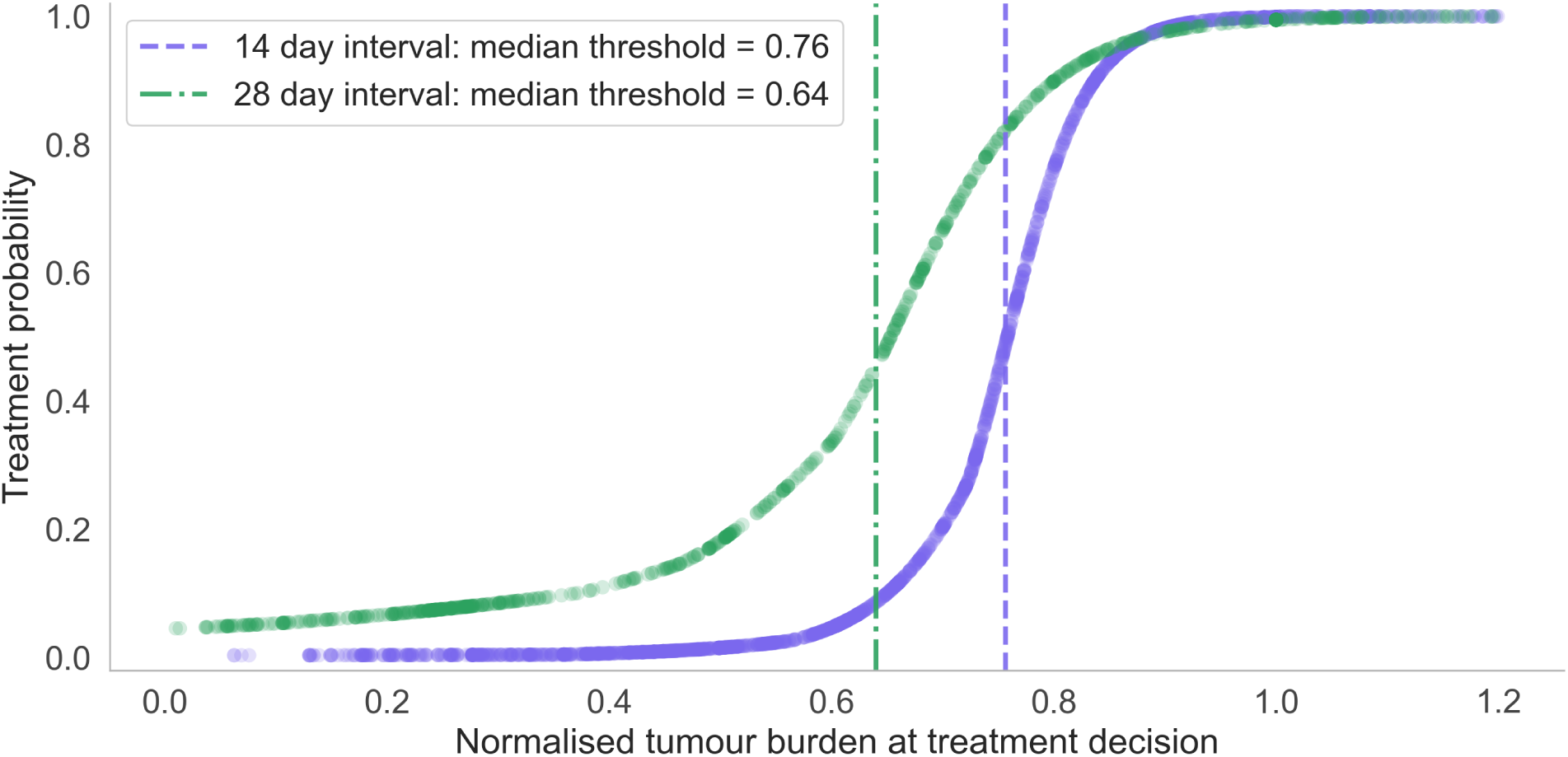
Population-level treatment-propensity boundaries. Treatment probability as a function of baseline-normalised tumour burden for the population-trained DRL policies at *d* = 14 and *d* = 28 days. The treatment-propensity boundary is defined as the tumour burden at which *p*_treat_ = 0.5. As observed for patient-specific DRL policies, the population-level boundary shifts to lower tumour-burden values as the decision interval increases.

Population-trained policies therefore provide a population-level treatment-propensity boundary that captures consistent trends in DRL policy structure and when applied improves TTP compared to MTD for most held-out patients, offering a practical starting point for treatment design. For example, at a clinically relevant decision interval of 4 weeks, the population-level upper threshold is 0.62 of the initial tumour burden. This suggests that under sparse, clinically realistic monitoring, delaying progression compared to MTD requires using a treatment-restart threshold lower than the initial tumour baseline. Unlike the Zhang et al. protocol which aims to keep tumour burden between two thresholds, the interpreted DRL boundary represents the learned probability of restarting treatment as tumour burden changes. The population-level boundary at *d* = 28 was 0.62 of baseline tumour burden, suggesting that under sparse, clinically realistic monitoring, TTP may be maximised by restarting treatment before the tumour returns to baseline.

Together, these results show that in NSCLC, learned treatment-propensity boundaries vary substantially between patients but exhibit consistent, quantifiable trends across decision intervals. This behaviour is consistent with the interval-dependent performance and robustness patterns observed in subsections 3.3 and 3.2.

Although the treatment-propensity boundary provides an interpretable summary of the learned behaviour, the full DRL policy remains the trained neural-network policy described in Methods and is only approximated by a deterministic binary switching rule. This is most evident in the few cases where DRL performs slightly worse than MTD on TTP, where a small decrease in TTP of a few days is traded for an increase in robustness. In these cases, the learned policy may stop treatment once the expected benefit of continued dosing becomes minimal, despite the tumour burden exceeding the treatment-propensity boundary. This behaviour reflects the full policy logic and cannot be captured by a single threshold alone.

### 3.5 Quality-adjusted survival reveals preference-dependent protocol rankings

The policies evaluated in this section were trained using the baseline survival-oriented reward. Here, QAS is used as an evaluation metric to assess how the same treatment policies would be valued under different illustrative patient preference profiles. Direct QoL-aware training is explored separately through reward shaping in Section 2.7.3.

#### 3.5.1 QAS across all protocols and preference profiles

We first evaluate all protocols using quality-adjusted-survival (QAS) as defined in Section 2.7.2. Because QAS depends on the weights assigned to the QoL components, individual patient preferences can lead to different QAS scores for the same treatment protocol. Across the evaluated decision intervals, median QAS is highest for the DRL protocol, indicating that DRL achieves the greatest cumulative quality-adjusted survival overall (Figure 9A). The DRL and Zhang et al. protocols show distinct patterns across profiles and decision intervals: For the medication-adverse and toxicity-adverse profiles, both the DRL and Zhang et al. protocol have higher median QAS compared to MTD. This reflects the combined effect of reduced treatment burden and longer survival. For these profiles, despite often using more medication and therefore having lower daily QoL than under the Zhang et al. protocol, the DRL protocol achieved the highest total QAS for these profiles as its longer TTP contributed more to the total QAS.

**Figure 9:**
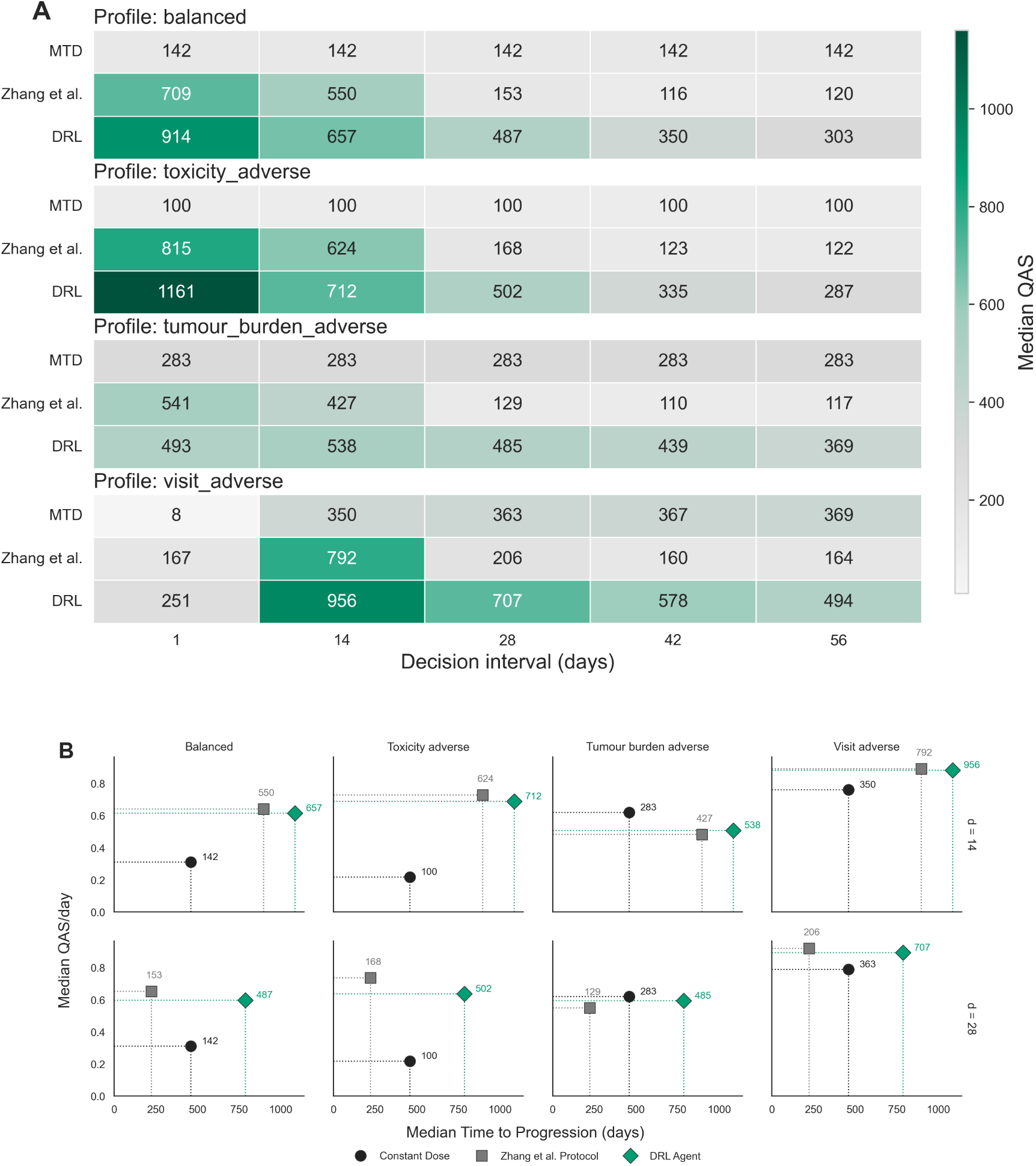
Quality-adjusted survival outcomes by preference profile and decision interval. (A) Heatmap of median quality-adjusted survival (QAS) by treatment protocol, preference profile, and decision interval. Rows represent treatment protocols, columns represent decision intervals (1, 14, 28, 42, and 56 days), and cell labels show median total QAS. (B) Median QAS/day versus median survival for selected decision intervals (14 and 28 days). Each point represents a treatment protocol within a preference profile, and point labels indicate the corresponding median total QAS. Together, the panels show how survival duration and daily quality of life contribute differently to total QAS across treatment protocols and decision intervals. **(a) Median QAS/day versus median survival for selected preference profiles at decision intervals of 14 and 28 days.** Each point represents one treatment protocol. The x-axis shows median survival (days), the y-axis shows median QAS/day, and adjacent numeric labels show the corresponding median total QAS.

For the balanced preference profile, the Zhang et al. protocol has high median QAS of 550 adjusted survival days at 14 day intervals, but by 42 day intervals performs worse than MTD due to the reduction in median TTP caused by premature progression. A similar pattern is observed for the tumour-burden-adverse profile across all decision intervals, however here the median daily QAS under MTD is much higher than for the other profiles, and the scores for the Zhang et al. and DRL protocols are lower. This result is consistent with the fact that evolutionary strategies tolerate and encourage higher tumour burdens, and are therefore penalised more strongly by this profile. These differences highlight how strongly patient preferences can change the perceived value of the same treatment protocol. The visit-adverse profile shows a distinct pattern. Because visit burden is fixed by the decision interval (*V* (*t*) = 1*/d*), differences in QAS under this weighting are driven mainly by survival rather than by changes in mean daily utility. This profile is therefore most informative for comparing QAS across decision intervals.

#### 3.5.2 DRL protocol gains in QAS arise mainly through longer survival

Total QAS combines survival duration with average QoL during survival. To separate these effects, we also examined the median QAS per day and plotted this against TTP for all protocols and preference profiles (Figure 9a). This analysis shows how the DRL advantage in total QAS is driven primarily by longer survival rather than consistently higher daily QoL. Although the medication-adverse and toxicity-adverse profiles represent distinct preferences (see Methods), they produced similar outcomes in the present setting because both medication burden and modelled toxicity are strongly linked to treatment exposure under binary ON/OFF dosing. To avoid visual redundancy, only the toxicity-adverse profile is shown. The distinction between these profiles may become more consequential in other settings with variable doses, alternative treatments, or patient-specific toxicity responses.

In the balanced and toxicity-adverse profiles, the Zhang et al. protocol often achieves a higher daily QoL score than DRL because it uses less medication. However, this higher daily QoL does not always translate into higher total QAS, particularly at longer decision intervals where the Zhang et al. protocol is more prone to premature progression. In contrast, DRL uses more medication as decision intervals increased, lowering daily QoL but increasing TTP (see supplementary materials S6).

For the tumour-burden-adverse profile, median QoL per day was highest under MTD as continuous treatment keeps tumour burden lower, however total QAS under MTD remained limited due to shorter TTP. This illustrates the trade-off inherent to this preference, keeping tumour burden low may improve daily quality of life but reduce cumulative QAS if it shortens TTP. This balancing act highlights the ecological dynamics of adaptive therapy. The DRL and Zhang et al. protocols deliberately tolerate a larger sensitive population to maintain competition and delay progression, however this is at odds with the tumour-burden-adverse preference profile.

Supplementary paired analyses showed that these QAS and preference profile interaction patterns also applied at the individual patient level. Across all preference profiles and decision intervals, the paired median ΔQAS for the DRL profile remained positive relative to MTD. Exact paired sign tests showed that this difference was significant across all profile and interval combinations. In contrast the QAS of the Zhang et al. protocol compared to MTD varied according to the preference profile and decision interval. The strongest example was for the tumour-burden-adverse profile where the Zhang et al. profile underperformed MTD at longer intervals. Full paired ΔQAS summaries, sign test results and patient level win rates for the four reported preference profiles are provided in Supplementary Tables S7–S9.

### 3.6 Reward shaping shifts survival-QoL trade-offs across profiles

Rather than directly incorporating QAS (see Section 2.7.2) into the reward function, we explored how different reward settings influence the DRL policy and QoL outcomes. To do this we performed a reward shaping sweep where we systematically varied the holiday threshold, holiday bonus, holiday penalty, and survival reward (Supplementary Section 2.6). For each reward configuration, a new DRL policy was trained and evaluated on two representative virtual patients.

Across the explored parameter ranges, the policy depended most on the relative weighting of the reward components rather than their exact values. In particular, a positive holiday bonus was important for encouraging treatment holidays. Without a reward associated with treatment holidays, the learned policy typically defaulted to MTD. The holiday threshold, i.e., the tumour burden threshold where treatment holidays switch from being rewarded to being punished, had the largest effect on treatment behaviour. Increasing the holiday threshold increased TTP as it allowed a higher tumour burden to be maintained. In contrast changes in the holiday bonus, holiday penalty, and survival reward had smaller effects on TTP.

The effect of reward shaping depends strongly on the preference profile. For the balanced and toxicity-adverse profiles, increasing TTP was not associated with a substantial reduction in median QAS. Several reward parameter configurations achieved both higher survival and high QAS compared to the other profiles. In these profiles, increasing TTP was not associated with a substantial reduction in median daily QoL. This suggests that for these preferences, the reward settings can be adjusted to prioritise survival without requiring a major decrease in daily QoL. In contrast, the tumour-burden-adverse profile showed a clear trade off, as reward settings associated with longer TTP generally produced lower daily QoL. These findings suggest that patient preferences can alter the degree to which evolutionary treatment principles should be applied. Patients who prioritise minimising tumour burden may prefer substantially lower holiday thresholds, accepting less ecological competition in exchange for tighter tumour control, whereas patients who prioritise toxicity reduction or treatment-free time may favour higher thresholds and longer treatment holidays.

Figure 10 illustrates how these reward-shaping trade-offs change the protocol and alter the tumour growth trajectory. Under the baseline reward, the DRL policy balances treatment holidays against tumour control, maintaining a substantial margin-to-failure while extending TTP relative to MTD. After increasing the holiday threshold, treatment holidays are rewarded at higher tumour burdens. This allows the sensitive population to remain dominant for longer, increasing the competitive suppression of resistant cells and extending TTP. However, the MTF has also decreased as the trajectory comes closer to the progression threshold. This demonstrates how the same reward modification that increases survival also reduces robustness.

**Figure 10:**
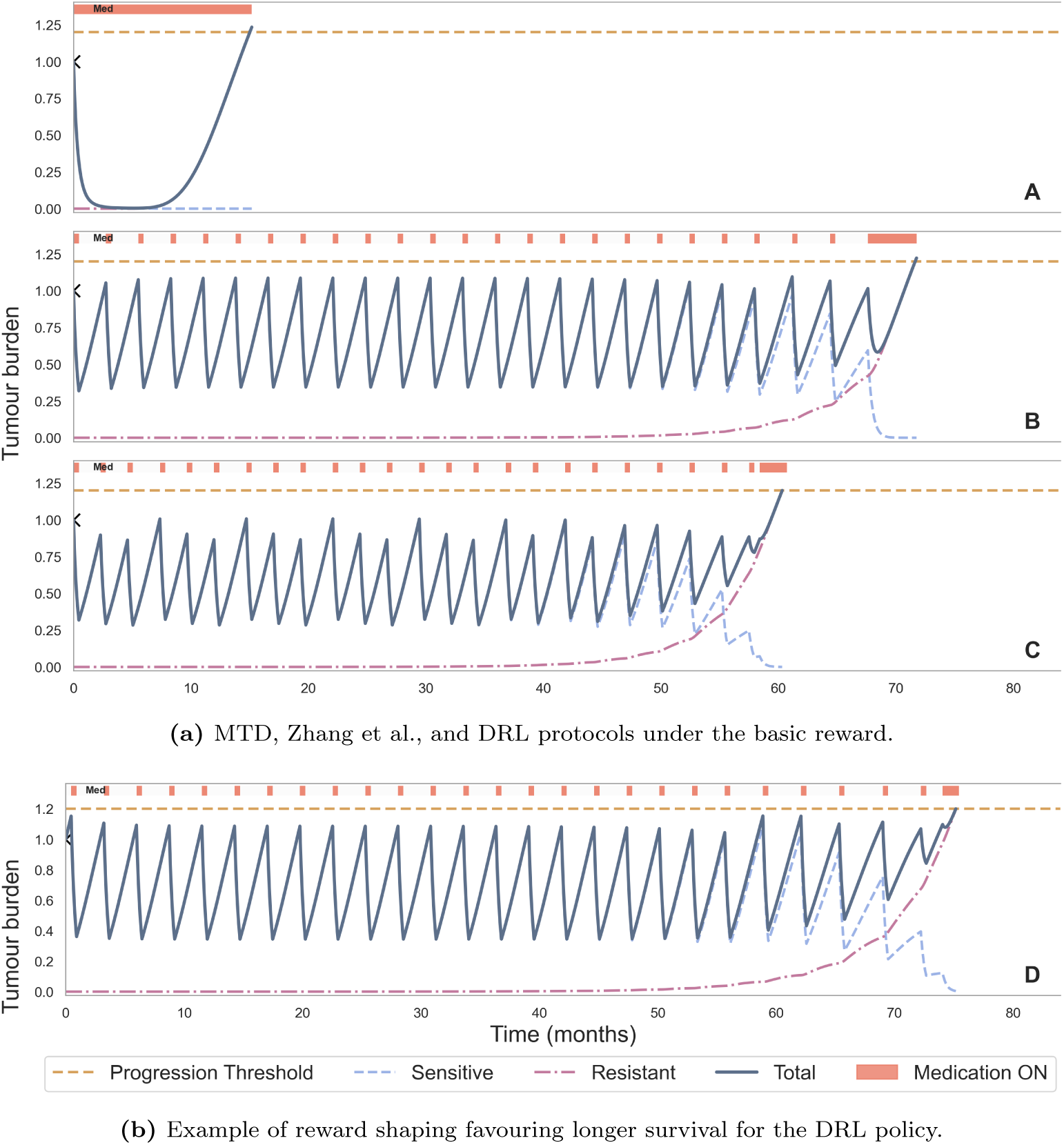
Trajectory-level illustration of the survival–robustness trade-off induced by reward shaping. Top: MTD, Zhang et al., and DRL protocols under the basic reward. Bottom: DRL policy after reward shaping to favour longer survival, shown here for patient 11 at decision interval *d* = 14 days with holiday threshold set to 1.2 and no holiday penalty. Increasing the holiday threshold prolongs time to progression, but reduces the margin to failure, illustrating that reward shaping can explicitly tune the balance between survival and safety.

## 4 Discussion

Despite strong theoretical foundations [10, 11, 37], adaptive cancer therapies remain challenging to implement clinically [28]. An adaptive protocol needs to include cancer type, medication, and individual patient tumour dynamics as well as accounting for the source and expected frequency of tumour burden measurements. DRL learned protocols fit naturally into this space as the policy is learned from interaction with a mechanistic environment rather than specified in advance. In this research we demonstrate the potential of DRL combined with mechanistic modelling to design flexible, robust and clinically feasible treatment protocols for non-small cell lung cancer. Flexibility means the policies are not predetermined, but adapt in response to both the individual patient’s tumour growth dynamics and to the time between observations. Robustness means the policies maintain a safety margin that prevents premature progression in cases where intended treatment restart is delayed. Clinical feasibility means that even at clinically realistic decision intervals the policies extend predicted time to progression compared to MTD. Beyond tumour control, our quality-adjusted-survival evaluation demonstrated how treatment experience varies with patient priorities, and our experiments on reward shaping demonstrate how learned treatment policies can be steered to align with a patient’s QoL preferences.

Previous work has shown that the Zhang et al. protocol can be vulnerable to premature progression arising from delayed treatment decisions and noisy observations [7, 29]. Our findings extend this by introducing a margin-to-failure metric to quantify robustness to delayed treatment restart. We then use this to show how DRL protocols maintain higher robustness by incorporating larger MTF, while still extending TTP compared to MTD. We show that because the fixed-threshold Zhang et al. protocol does not adapt to individual patient dynamics, its success is interval-dependent, with the interaction of decision interval and individual patient dynamics determining the success or failure of the protocol, and more cases of premature progression as the interval increased. In contrast the DRL agent adapted to increasing decision intervals by lowering the containment threshold. This is a learned response to the uncertainty of outcomes when decisions are far apart, i.e. when decisions are infrequent, maintaining a high tumour burden becomes more risky as tumour growth between observations is more likely to lead to premature progression.

Improved robustness under sparse monitoring is not obtained without cost. As decision intervals increased, DRL policies used medication more frequently and in some cases approached continuous MTD, indicating that robustness to delayed or infrequent decisions is partly achieved by sacrificing some of the medication sparing that motivates evolutionary therapy. Our reward shaping experiments made this trade-off explicit, showing that increasing the holiday threshold could prolong time to progression, but at the cost of a reduced margin-to-failure. Interestingly, the same holiday threshold parameter which produced the largest changes in TTP also had the strongest influence on robustness, indicating that threshold selection is the most important reward component governing the balance between tumour control and premature progression risk. This is biologically interpretable, because the holiday threshold directly determines the size of the surviving sensitive population. Higher thresholds preserve more ecological competition and therefore suppress resistant clones more effectively, but they also leave the system closer to the progression boundary. Lower thresholds sacrifice some competitive suppression in exchange for greater robustness against delayed treatment restart. Our findings are consistent with previous work on population-trained policies for prostate cancer which showed that population-trained policies achieved greater robustness to inter-patient variability by learning lower thresholds than were seen with most the individually-trained policies [6]. While this approach was robust to different patient dynamics and increased TTP relative to MTD for the majority of patients, the robustness involved a trade-off with tumour control as TTP was reduced compared to the individual policies. Similar results were seen under the population-trained policy in our study, where at 28 day decision intervals, the threshold was 62% of the initial tumour burden, and 90% of previously unseen virtual patients showed increased TTP compared to MTD.

This generalisation matters more in NSCLC than in prostate cancer. In NSCLC, tumour burden is typically assessed using imaging at intervals several weeks apart. Combined with rapid tumour growth, this leaves little time and limited data with which to fit patient-specific mechanistic models. This differs from the slower growing prostate cancer where PSA provides frequent more frequent measurements over a longer time period. This means that while creating an individual treatment schedule by updating a pre-trained population model may be feasible in prostate cancer [6], this approach is unlikely to work in NSCLC. In this context, using a population-trained protocol directly may be an alternative.

The upper containment thresholds we found were notably lower than those used in previous applications of adaptive therapy or in theoretically optimal perfect monitoring conditions. For example, using 62% of the initial tumour burden at four weekly intervals. In the NSCLC setting, this may offer a more practical and potentially safer starting point than applying the Zhang et al. protocol, especially given the substantial rate of premature progression we observed in simulations of the Zhang et al. protocol under longer decision intervals. In aggressive cancers such as NSCLC, a lower restart threshold may offer a safer option as it preserves a margin to progression while still allowing treatment holidays to control the growth of resistance. The learned DRL policies balance tumour burden with maintaining sufficient sensitive cells to delay the growth of resistance through competition. Monitoring frequency shifts this balance, as maintaining a large sensitive population becomes riskier when tumour growth is observed infrequently. It has been previously shown that the ideal tumour containment strategy can delay resistance longer by keeping the total tumour volume as close as possible to an upper threshold [37], however this ideal assumes continuous monitoring which the authors acknowledge is unfeasible. Recent work has also explored single-threshold adaptive therapy using partial treatment during fixed decision intervals [19]. In our setting DRL protocols can be interpreted as a similar strategy, with a single upper treatment-propensity bound [7]. Under daily observations *d* = 1, the learned policies provided high thresholds close to and even above the initial tumour burden. However, as the decision intervals increase and observations become sparser the containment becomes progressively lower, and the predicted TTP becomes shorter though remaining above MTD. Importantly, the learned policies were not just simple fixed thresholds, but varied according to the individual tumour dynamics and decision interval. At longer intervals the DRL policy recommended MTD for some patients, effectively a containment threshold of 0 where treatment is always on. In other cases, when competitive release had already occurred treatment was stopped even if the tumour burden had passed the threshold, indicating a more complex policy that balances treatment holiday with survival benefit.

Previous work combining reinforcement learning with mechanistic modelling to develop adaptive treatment protocols has focused on time to progression as a single outcome metric [6, 13, 17, 18, 34]. In contrast, our study evaluates treatment strategies using tumour control, robustness, and QoL metrics. By measuring quality-adjusted-survival outcomes, we show how patient preference changes the value of a treatment protocol. Then, by mapping how QoL objectives can be incorporated through reward shaping, we demonstrate how DRL can expand treatment protocol design beyond tumour control to incorporate patient preferences.

The policies compared under QAS were trained with the baseline survival-oriented reward, so this analysis asks how the same protocols would be valued under different patient preferences rather than how policies optimised for those preferences would behave. Within this framing, our results indicate that adaptive therapy should be judged not only by tumour control, but also by how different patients value the experience of treatment. In the medication-adverse and toxicity-adverse preference profiles the lower medication use under the DRL and Zhang et al. protocols improved daily QoL, however for the tumour-burden-adverse profile there was a clear trade-off between keeping tumour burden low and extending survival. Although total QAS under DRL remained higher than under MTD for this profile, the gain came from longer TTP, and daily QoL was lower than under MTD. In practice, this means that QoL-aware treatment protocols must balance competing objectives including survival, robustness, toxicity, and treatment burden. This is consistent with other work on multi-objective optimisation for cancer treatment where treatment design is framed as a process of navigating trade-offs between often conflicting goals [36]. This is consistent with the limited literature on QoL-aware adaptive therapy. For example, Gevertz et al. showed that incorporating toxicity constraints can extend time to progression [12]; our results similarly suggest that reducing toxicity and medication burden can align well with extending TTP using adaptive therapy approaches, whereas maintaining low tumour burden involves a more direct trade-off with survival. In effect, for some preference profiles there is a trade-off between “living longer” and “living better per day” and the right treatment decision depends on the patient.

Several limitations should be considered. First, the tumour environment is based on a simplified two-population ODE model and does not include aspects of tumour biology such as spatial structure or potential phenotypic plasticity. The action space is binary (treatment ON/OFF), which limits how finely different QoL objectives can be separated. It also means there is no possibility for using lower doses which may control tumour burden and improve QoL. In addition, the QoL utility used here is a proxy derived from tumour burden, toxicity, medication exposure, and monitoring burden rather than from observed patient-reported outcomes, and the reward-shaping sweep was illustrative rather than a population-level optimisation study. The margin-to-failure metric provides a practical way to quantify safety against delayed restart or modest tumour-burden misestimation. However, because measurement error was not modelled explicitly, future work should evaluate learned policies under stochastic imaging noise and biased tumour-burden estimates to determine how well MTF predicts real-world robustness. Our QoL formulation also assumes that treatment toxicity depends on treatment exposure and tumour burden, but does not explicitly distinguish the clinical impact of sensitive and resistant subpopulations. Future work could incorporate composition-dependent QoL terms, allowing resistant disease or higher resistance levels to contribute differently to symptoms, metastatic risk, or treatment burden, as implemented in a recent study where patient quality-of-life depends on tumour burden, treatment dose, and treatment-induced resistance [27].

These limitations also point to important future directions. Extending the action space to include multiple dose levels, or even multiple medications, would greatly expand the range of clinically meaningful adaptive strategies and would also make QoL extensions more expressive. Richer action spaces would allow medication burden, toxicity, and tumour control to be separated more clearly than in the current binary setting. A second important extension would be to optimise decision windows in addition to treatment actions. In the present framework, visit burden was included in QoL evaluation, but because the monitoring interval was fixed externally it was not itself directly controllable by the policy. Allowing the agent to optimise when reassessment occurs would better accommodate visit-adverse preferences and make monitoring burden a true decision variable. The same framework could also be extended to include cost-aware objectives combining medication exposure and visit frequency. Future extensions could also incorporate broader treatment-burden dimensions, including financial toxicity, caregiver burden, and patient-reported outcomes, which were not explicitly modelled here.

In summary, our findings show that monitoring frequency determines how safely competition between drug-sensitive and drug-resistant cells can be exploited during evolutionary therapy. DRL adapted this balance to the treatment decision intervals, by restarting treatment at lower tumour burdens as monitoring became less frequent and by approaching continuous treatment when safe treatment holidays could not be identified. These results suggest that evolutionary therapy should not rely on single predetermined containment thresholds across decision intervals, but should account for the interaction between tumour growth, competitive dynamics and the available monitoring schedule. Although prospective validation is required, this study provides a biologically informed framework for developing robust and patient-centred evolutionary therapies for EGFR-mutated NSCLC and other fast-growing cancers.

## Data Availability

Virtual patient cohort data, parameter distributions, and the code used to generate virtual patients, train the DRL agents, and reproduce all analyses are available at https://gitlab.tudelft.nl/evolutionary-game-theory-lab/drl_for_qol_aware_nsclc. The virtual patients were generated from parameter distributions derived from a model fitted to START-TKI data (as reported in Jansén-Storbacka et al. [16]). The underlying START-TKI individual patient data are not publicly available because of privacy and ethical restrictions but may be made available upon reasonable request to Anne-Marie Dingemans at Erasmus Medical Center, subject to institutional and ethical approvals [5].

## References

[1] Freddie Bray, Mathieu Laversanne, Hyuna Sung, Jacques Ferlay, Rebecca L. Siegel, Isabelle Soerjomataram, and Ahmedin Jemal. 2024. Global cancer statistics 2022: GLOBOCAN estimates of incidence and mortality worldwide for 36 cancers in 185 countries. CA: A Cancer Journal for Clinicians 74, 3 (2024), 229–263. doi:10.3322/caac.21834

[2] Juliann Chmielecki, Jasmine Foo, Geoffrey R. Oxnard, Katherine Hutchinson, Kadoaki Ohashi, Romel Somwar, Lu Wang, Katherine R. Amato, Maria Arcila, Martin L. Sos, Nicholas D. Socci, Agnes Viale, Elisa De Stanchina, Michelle S. Ginsberg, Roman K. Thomas, Mark G. Kris, Akira Inoue, Marc Ladanyi, Vincent A. Miller, Franziska Michor, and William Pao. 2011. Optimization of dosing for EGFR-mutant non-small cell lung cancer with evolutionary cancer modeling. Science Translational Medicine 3, 90 (July 2011), 90ra59. doi:10.1126/scitranslmed.3002356

[3] Antoine M Dujon, Athena Aktipis, Catherine Alix-Panabières, Sarah R Amend, Amy M Boddy, Joel S Brown, Jean-Pascal Capp, James DeGregori, Paul Ewald, Robert A Gatenby, Marco Gerlinger, Mathieu Giraudeau, Rodrigo K Hamede, Elsa Hansen, Irina Kareva, Carlo C Maley, Andriy Marusyk, Nicholas McGranahan, Michael J Metzger, Aurora M Nedelcu, Robert Noble, Leonard Nunney, Kenneth J Pienta, Kornelia Polyak, Pascal Pujol, Andrew F Read, Benjamin Roche, Susanne Sebens, Eric Solary, Holly Staňková, Kateřinaand Swain Ewald, Frédéric Thomas, and Beata Ujvari. 2021. Identifying key questions in the ecology and evolution of cancer. Evolutionary Applications 14, 4 (2021), 877–892. doi:10.1111/eva.13190

[4] E A Eisenhauer, P Therasse, J Bogaerts, L H Schwartz, D Sargent, R Ford, J Dancey, S Arbuck, S Gwyther, M Mooney, L Rubinstein, L Shankar, L Dodd, R Kaplan, D Lacombe, and J Verweij. 2009. New response evaluation criteria in solid tumours: revised RECIST guideline (version 1.1). European Journal of Cancer (Oxford, England: 1990) 45, 2 (1 2009), 228–247. doi:10.1016/j.ejca.2008.10.026

[5] Erasmus Medical Center. 2022. proSpecTive sAmpling in dRiver muTation Pulmonary Oncology Patients on Tyrosine Kinase Inhibitors (START-TKI). https://clinicaltrials.gov/study/NCT05221372. https://clinicaltrials.gov/study/NCT05221372 Observational prospective cohort study in NSCLC patients with oncogenic driver mutations treated with tyrosine kinase inhibitors. Estimated enrollment: 1300 participants. Study start: 2017-02-02. Estimated completion: 2031-01-01. Principal Investigator: Anne-Marie Dingemans, MD, PhD..

[6] Kit Gallagher, Maximilian A.R. Strobl, Derek S. Park, Fabian C. Spoendlin, Robert A. Gatenby, Philip K. Maini, and Alexander R.A. Anderson. 2024. Mathematical Model-Driven Deep Learning Enables Personalized Adaptive Therapy. Cancer Research 84, 11 (2024), 1929–1941. doi:10.1158/0008-5472.CAN-23-2040

[7] Kit Gallagher, Maximilian A. R. Strobl, Alexander R. A. Anderson, and Philip K. Maini. 2025. Deriving Optimal Treatment Timing for Adaptive Therapy: Matching the Model to the Tumor Dynamics. Bulletin of Mathematical Biology 87, 146, Article 146 (2025), 19 pages. doi:10.1007/s11538-025-01525-y

[8] Kit Gallagher, Maximilian A. R. Strobl, Derek S. Park, Fabian C. Spoendlin, Robert A. Gatenby, Philip K. Maini, and Alexander R. A. Anderson. 2024. Supplementary Information for: Mathematical Model-Driven Deep Learning Enables Personalized Adaptive Therapy. Supplementary material. https://aacrjournals.org/cancerres/article/84/11/1929/745515/Mathematical-Model-Driven-Deep-Learning-Enables

[9] Robert A. Gatenby. 2009. A Change of Strategy in the War on Cancer. Nature 459, 7246 (May 2009), 508–509. doi:10.1038/459508a

[10] Robert A Gatenby and Joel S Brown. 2018. The evolution and ecology of resistance in cancer therapy. doi:10.1101/cshperspect.a033415

[11] Robert A. Gatenby, Ariosto S. Silva, Robert J. Gillies, and B. Roy Frieden. 2009. Adaptive therapy. Cancer Research 69, 11 (6 2009), 4894–4903. doi:10.1158/0008-5472.CAN-08-3658

[12] Jana L. Gevertz, Harsh Vardhan Jain, Irina Kareva, Kathleen P. Wilkie, Joel Brown, Yitong Pepper Huang, Eduardo Sontag, Vladimir Vinogradov, and Mark Davies. 2026. Delaying cancer progression by integrating toxicity constraints in a model of adaptive therapy. npj Systems Biology and Applications 12, 1 (2026), 11. doi:10.1038/s41540-025-00635-6

[13] M. Giles and P. K. Newton. 2025. Chemotherapy dose scheduling via Q-learning in a Markov tumor model. bioRxiv (June 2025). doi:10.1101/2025.06.05.658118 preprint.

[14] Lizza E. L. Hendriks, Anne-Marie C. Dingemans, Dirk K. M. De Ruysscher, Mieke J. Aarts, Lidia Barberio, Robin Cornelissen, Koen J. Hartemink, Michel van den Heuvel, Ed Schuuring, Hans J. M. Smit, Antonie J. van der Wekken, and Egbert F. Smit. 2021. Lung Cancer in the Netherlands. Journal of Thoracic Oncology 16, 3 (2021), 355–365. doi:10.1016/j.jtho.2020.10.012

[15] Roy S. Herbst, Daniel Morgensztern, and Chris Boshoff. 2018. The biology and management of non-small cell lung cancer. Nature 553, 7689 (Jan. 2018), 446–454. doi:10.1038/nature25183

[16] Laura R. Jansén-Storbacka, Kailas S. Honasoge, Eva Molnárová, Arina Soboleva, Bram C. Agema, Marthe S. Paats, Dirk Jan A. R. Moes, G. D. Marijn Veerman, Alethea B. T. Barbaro, Roel Dobbe, Irene Grossmann, Sepinoud Azimi, Ron H. J. Mathijssen, Anne-Marie C. Dingemans, and Kateřina Staňková. 2026. Can evolutionary therapy be applied in non-small cell lung cancer? Scientific Reports 16, 1 (2026), 36712. doi:10.1038/s41598-026-36712-x

[17] Zhiqing Li, Yun Zhao, Zhiqiang Yu, and Xuewen Tan. 2026. Adaptive therapy for non-small cell lung cancer via integrated Stackelberg game and deep reinforcement learning frameworks. AIP Advances 16, 1 (2026), 015113. doi:10.1063/5.0306241

[18] Yitao Lu, Qian Chu, Zhen Li, Mengdi Wang, Robert Gatenby, and Qingpeng Zhang. 2024. Deep Reinforcement Learning Identifies Personalized Intermittent Androgen Deprivation Therapy for Prostate Cancer. Briefings in Bioinformatics 25, 2 (2024), bbae071. doi:10.1093/bib/bbae071

[19] Kexin Ma, Ningjing Wang, Zai Yang, Robert A. Cheke, and Biao Tang. 2026. Single-threshold-guided adaptive cancer therapy with partial-cycle treatment: A mechanistic and reinforcement learning analysis. PLOS Computational Biology 22, 6 (2026), e1014457. doi:10.1371/journal.pcbi.1014457

[20] Andriy Marusyk, Vanessa Almendro, and Kornelia Polyak. 2012. Intra-tumour heterogeneity: a looking glass for cancer? Nature Reviews Cancer 12, 5 (2012), 323–334. doi:10.1038/nrc3261

[21] Lauren M F Merlo, John W Pepper, Brian J Reid, and Carlo C Maley. 2006. Cancer as an evolutionary and ecological process. Nature Reviews Cancer 6, 12 (12 2006), 924–935. doi:10.1038/nrc2013

[22] A. C. Mita, K. Papadopoulos, M. J. A. de Jonge, G. Schwartz, J. Verweij, M. M. Mita, A. Ricart, Q. S.-C. Chu, A. W. Tolcher, L. Wood, S. McCarthy, M. Hamilton, K. Iwata, B. Wacker, K. Witt, and E. K. Rowinsky. 2011. Erlotinib ‘dosing-to-rash’: a phase II intrapatient dose escalation and pharmacologic study of erlotinib in previously treated advanced non-small cell lung cancer. British Journal of Cancer 105, 7 (2011), 938–944. doi:10.1038/bjc.2011.332

[23] Eva Molnárová, Ties A. Mulders, Marcela Spee-Dropková, Louise M. Spekking, Sepinoud Azimi, Irene Grossmann, Anne-Marie C. Dingemans, and Kateřina Staňková. 2026. Enabling Evolutionary Therapy in Metastatic Cancer Lacking Serum Biomarkers. Journal of Evolutionary Biology (2026). arXiv:2512.19485 [q-bio.PE] doi:10.48550/arXiv.2512.19485 In press.

[24] David G. Moriarty, Mathew M. Zack, and Rosemarie Kobau. 2003. The Centers for Disease Control and Prevention’s Healthy Days Measures - Population tracking of perceived physical and mental health over time. Health and Quality of Life Outcomes 1, 1 (2003), 37. doi:10.1186/1477-7525-1-37

[25] Kenneth J Pienta, Emma U Hammarlund, Robert Axelrod, Sarah R Amend, and Joel S Brown. 2020. Convergent evolution, evolving evolvability, and the origins of lethal cancer. Molecular Cancer Research 18, 6 (2020), 801–810. doi:10.1158/1541-7786.MCR-19-1158

[26] Charles M. Rudin, Wanqing Liu, Apurva Desai, et al. 2008. Pharmacogenomic and pharmacokinetic determinants of erlotinib toxicity. Journal of Clinical Oncology 26, 7 (2008), 1119–1127. doi:10.1200/JCO.2007.13.1128

[27] Monica Salvioli, Hasti Garjani, Mohammadreza Satouri, Mark Broom, Yannick Viossat, Joel S Brown, Johan Dubbeldam, and Kateřina Staňková. 2024. Stackelberg evolutionary games of cancer treatment: What treatment strategy to choose if cancer can be stabilized? Dynamic Games and Applications 14, 1 (2024), 1–20. doi:10.1007/s13235-024-00609-z

[28] Arina Soboleva, Irene Grossmann, Anne-Marie C. Dingemans, Jafar Rezaei, and Kateřina Staňková. 2025. Bringing evolutionary cancer therapy to the clinic: a systems approach. npj Systems Biology and Applications 11, 1 (2025), 56. doi:10.1038/s41540-025-00528-8

[29] Arina Soboleva, Kailas S. Honasoge, Eva Molnárová, Ties A. Mulders, Anne-Marie C. Dingemans, Irene Grossmann, Jafar Rezaei, and Kateřina Staňková. 2026. Assessing the Operational Feasibility of Evolutionary Therapy in Metastatic Non-Small Cell Lung Cancer. bioRxiv (2026). doi:10.64898/2026.02.25.707957 Preprint; version 1 posted 26 Feb 2026.

[30] Kateřina Staňková. 2019. Resistance games. Nature Ecology & Evolution 3, 3 (3 2019), 336–337. doi:10.1038/s41559-018-0785-y

[31] Kateřina Staňková, Joel S. Brown, William S. Dalton, and Robert A. Gatenby. 2019. Optimizing Cancer Treatment Using Game Theory: A Review. JAMA Oncology 5, 1 (1 2019), 96–103. doi:10.1001/jamaoncol.2018.3395

[32] Christi M.J. Steendam, G. D. Marijn Veerman, Melinda A. Pruis, Peggy Atmodimedjo, Marthe S. Paats, Cor van der Leest, Jan H. von der Thüsen, David C.Y. Yick, Esther Oomen De Hoop, Stijn L.W. Koolen, Winand N.M. Dinjens, Ron H.N. van Schaik, Ron H.J. Mathijssen, Joachim G.J.V. Aerts, Hendrikus Jan Dubbink, and Anne Marie C. Dingemans. 2020. Plasma predictive features in treating egfr-mutated non-small cell lung cancer. Cancers 12, 11 (11 2020), 1–17. doi:10.3390/cancers12113179

[33] Alexander Stein, Monica Salvioli, Hasti Garjani, Johan Dubbeldam, Yannick Viossat, Joel S. Brown, and Kateřina Staňková. 2023. Stackelberg evolutionary game theory: how to manage evolving systems. Philosophical Transactions of the Royal Society B: Biological Sciences 378, 1876 (2023), 20210495. doi:10.1098/rstb.2021.0495

[34] F. Tavakoli, D. Mohammadpur, J. S. Sartakhti, and M. H. Manshaei. 2025. Reinforcement Learning-Driven Evolutionary Stackelberg Game Model for Adaptive Breast Cancer Therapy. Mathematical and Computational Applications 30, 6 (2025), 134. doi:10.3390/mca30060134

[35] The WHOQOL Group. 1995. The World Health Organization Quality of Life Assessment (WHOQOL): Position paper from the World Health Organization. Social Science & Medicine 41, 10 (1995), 1403–1409. doi:10.1016/0277-9536(95)00112-K

[36] Maicon de Paiva Torres, Fran Sérgio Lobato, and Gustavo Barbosa Libotte. 2025. Exploring trade-offs in drug administration for cancer treatment: A multi-criteria optimisation approach. Mathematical Biosciences 382 (2025), 109404. doi:10.1016/j.mbs.2025.109404

[37] Yannick Viossat and Robert Noble. 2021. A theoretical analysis of tumour containment. Nature Ecology and Evolution 5, 6 (6 2021), 826–835. doi:10.1038/s41559-021-01428-w

[38] Jung-Der Wang and Jing-Shiang Hwang. 2014. Quality-Adjusted Survival. Springer Netherlands, Dordrecht, 5322–5325. doi:10.1007/978-94-007-0753-5_2388

[39] Jeffrey West, Fred Adler, Jill Gallaher, Maximilian Strobl, Renée Brady-Nicholls, Joel Brown, Mark Roberson-Tessi, Eunjung Kim, Robert Noble, Yannick Viossat, David Basanta, and Alexander R.A. Anderson. 2023. A survey of open questions in adaptive therapy: Bridging mathematics and clinical translation. doi:10.7554/elife.84263

[40] Jeffrey West, Li You, Jingsong Zhang, Robert A. Gatenby, Joel S. Brown, Paul K. Newton, and Alexander R.A. Anderson. 2020. Towards multidrug adaptive therapy. Cancer Research 80, 7 (4 2020), 1578–1589. doi:10.1158/0008-5472.CAN-19-2669

[41] Benjamin Wölfl, Hedy te Rietmole, Monica Salvioli, Artem Kaznatcheev, Frank Thuijsman, Joel S. Brown, Boudewijn Burgering, and Kateřina Staňková. 2022. The Contribution of Evolutionary Game Theory to Understanding and Treating Cancer. Dynamic Games and Applications 12, 2 (2022), 313–342. doi:10.1007/s13235-021-00397-w

[42] Jingsong Zhang, Jessica Cunningham, Joel S. Brown, and Robert A. Gatenby. 2022. Evolution-based mathematical models significantly prolong response to abiraterone in metastatic castrate-resistant prostate cancer and identify strategies to further improve outcomes. eLife 11 (June 2022), e76284. doi:10.7554/eLife.76284

[43] Jingsong Zhang, Jessica J. Cunningham, Joel S. Brown, and Robert A. Gatenby. 2017. Integrating Evolutionary Dynamics into Treatment of Metastatic Castrate-Resistant Prostate Cancer. Nature Communications 8 (2017), 1816. doi:10.1038/s41467-017-01968-5

[44] Jingsong Zhang, Jill Gallaher, Liang Wang, Monica Sheila Chatwal, Youngchul Kim, and Robert A Gatenby. 2022. A phase 1b adaptive androgen deprivation therapy trial in metastatic castration sensitive prostate cancer. Journal of Clinical Oncology 40, 16 suppl (6 2022), 5075. doi:10.1200/JCO.2022.40.16{_}suppl.5075

