## Supplementary methods for "Robust and Quality-of-Life-Aware Treatment Protocols in NSCLC using Deep Reinforcement Learning"

[†]Joint last authors

### 1 Virtual Patient Generation

#### 1.1 Overview and data source

Virtual patients were generated by sampling parameter sets $(r_S, r_R, K, \lambda, \alpha_{SR}, \alpha_{RS}, x_S(0), x_R(0))$ for the model in Equation 1. Parameter means and variances for the growth rates, carrying capacity, drug effect, and initial populations were taken from our previous fits to NSCLC patients treated with erlotinib [8]. The full sampling distributions, parameter values, and truncation bounds are reported in Table S1.

#### 1.2 Parameter distributions and priors

Parameters were sampled independently from distributions informed by previous individual fits. We did not enforce correlations between fitted parameters as correlations estimated from a small population may artificially restrict the diversity of the virtual patients. Independent sampling of parameters was therefore used to allow the protocols to be tested across a broader range of tumour dynamics. For log-normally sampled parameters, the fitted means and variances were used to define the underlying distributions, and samples were drawn after truncation to the bounds in Table S1. The realised sampled moments therefore need not exactly match the fitted moments. For the initial sensitive and resistant populations, the means and standard deviations reported in Table S1 correspond to the fitted values. During virtual-patient generation, the fitted variances were doubled (equivalently, the standard deviations were multiplied by $\sqrt{2}$) before constructing the log-normal sampling distributions to increase heterogeneity in baseline tumour size and composition.

Although our previous NSCLC modelling produced point estimates for the competition coefficients $\alpha_{SR}$, and $\alpha_{RS}$, repeated fits for the same patient sometimes yielded varying parameter values despite nearly identical tumour trajectories, indicating practical non-identifiability. This is consistent with evidence that competition coefficients are harder to estimate than demographic rates [7, 11]. We therefore sampled the competition coefficients from a uniform distribution $\alpha_{SR}, \alpha_{RS} \sim \mathcal{U}(0.1, 2)$. This range spans weak to strong competition and is consistent with previous work showing that these ranges permit coexistence of sensitive and resistant populations in previous modelling studies [2].

| **Parameter** | **Description** | **Units** | **Distribution** | **Bounds** |
|---|---|---|---|---|
| $r_S$ | Sensitive growth rate | day$^{-1}$ | LogNormal, $m = 0.042$, $s = 0.035$ | $[0.005,\ 0.1]$ |
| $r_R$ | Resistant growth rate | day$^{-1}$ | LogNormal, $m = 0.018$, $s = 0.025$ | $[0.005,\ 0.1]$ |
| $\lambda$ | Effect of medication | mg$^{-1}$day$^{-1}$ | LogNormal, $m = 5.73 \times 10^{-4}$, $s = 2.64 \times 10^{-4}$ | $[10^{-5},\ 2 \times 10^{-3}]$ |
| $K$ | Carrying capacity | mm$^3$ | LogNormal, $m = 6.94 \times 10^4$, $s = 3.85 \times 10^4$ | $[2 \times 10^4,\ 3 \times 10^5]$ |
| $\alpha_{SR}$ | Competition: R → S | dimensionless | $\mathcal{U}(0.1, 2)$ | $[0.1,\ 2]$ |
| $\alpha_{RS}$ | Competition: S → R | dimensionless | $\mathcal{U}(0.1, 2)$ | $[0.1,\ 2]$ |
| **Initial conditions** | Initial sensitive burden | mm$^3$ | LogNormal, $m = 4.3 \times 10^4$, $s = 1.8 \times 10^4$ | $[10^4,\ 10^5]$ |
| | Initial resistant burden | mm$^3$ | LogNormal, $m = 0.215$, $s = 0.135$ | $[0.005,\ 1]$ |

Table S1: **Parameter distributions and bounds used for virtual patient generation.** The parameters $r_S, r_R, \lambda$ and $K$ were sampled from truncated log-normal distributions based on previous individual fits to patient data [8]. Sampling constraints were used to ensure biologically plausible growth dynamics and carrying capacities.

#### 1.3 Sampling procedure

To improve sampling efficiency, parameters were generated sequentially and constraints such as $r_S > r_R$ were enforced directly during sampling. For example, after sampling $r_S$, the resistant growth rate $r_R$ was sampled from its truncated distribution with upper bound $r_S$. Similarly, carrying capacities were sampled only from ranges compatible with the sampled initial tumour burden. This approach ensured that biologically implausible parameter combinations were excluded during sampling rather than through repeated rejection, substantially improving acceptance rates.

Sampling constraints were applied during virtual-patient generation to ensure biologically plausible parameter sets. We enforced $r_S > r_R$, constrained carrying capacities to satisfy

$$1.5\big(x_S(0) + x_R(0)\big) < K < 5\big(x_S(0) + x_R(0)\big),$$

and required the medication effect $\lambda$ to be sufficiently large relative to $r_S$ so that treatment at the full 150 mg dose could induce a net decline in the sensitive population when medication was ON.

##### 1.4 Trajectory screening

After parameter sampling, each virtual patient was simulated under continuous MTD to identify clinically realistic tumour trajectories. Sampled parameter sets were retained only if they exhibited an initial treatment response followed by a 20% increase in spherical-equivalent tumour diameter within two years. These trajectory-level filters were used to exclude unrealistic tumour dynamics that would not be expected in EGFR-mutant NSCLC treated with erlotinib. For cohort filtering, tumour volume was converted to a spherical-equivalent diameter so that RECIST-like diameter criteria could be applied.

##### 1.5 Cohort size and reproducibility

All virtual-patient sampling and simulations used NumPy's random number generator. Sampling continued until 60 virtual patients were selected. In this run, 60 were accepted from 150 attempts (40% acceptance). The accepted cohort (60 patients) is provided as a CSV file to ensure exact reproducibility. Table S1 lists the log-normal distributions and the truncation bounds used for sampling virtual patient parameters.

##### 1.6 Practical non-identifiability of parameters

Tumour growth models based on ordinary differential equations, which typically form the environment for these reinforcement learning systems, may suffer from practical nonidentifiability of parameters when fitted to sparse clinical time series [3]. When only a few measurements are available for an individual patient, many different parameter sets can reproduce the observed tumour trajectory with comparable quality, so repeated fittings usually generate a broad distribution of plausible values. This distribution reflects epistemic uncertainty arising from limited clinical information and from simplifications inherent in

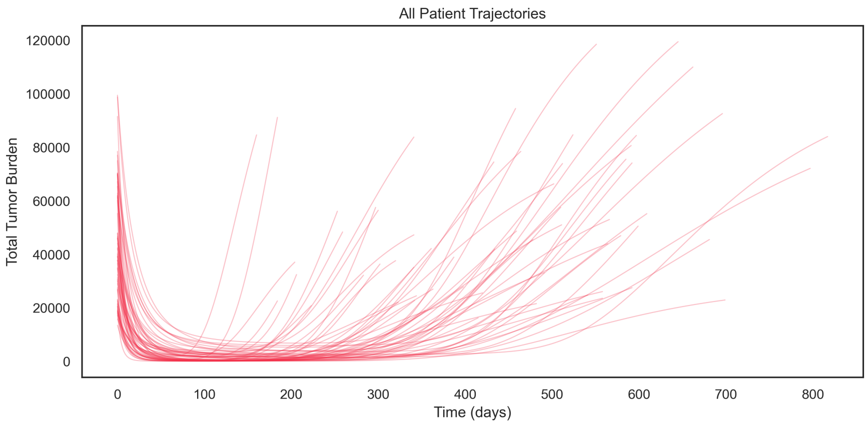


Figure S1: Variation in tumour initial conditions and growth for the virtual patient cohort.

the model, in addition to the aleatoric uncertainty introduced by measurement noise. Even when a model is structurally identifiable in principle, practical nonidentifiability under real clinical conditions can lead to misleading biological interpretations and unreliable predictions, and reliance on single point estimates in such circumstances has been discouraged [3]. More generally, it has been argued that mathematical oncology models should be rigorously parametrised, calibrated, and validated before being used to guide treatment decisions, since insufficient data and nonidentifiable parameters can result in unreliable therapeutic recommendations [1]. In reinforcement learning settings, if the agent is trained using only one fitted value for each parameter, any bias or instability in these estimates may be inherited by the learned policy, potentially degrading performance when the true patient parameters differ from those used in training.

#### 1.7 Log-normal sampling

Log-normal distributions are widely observed across biological and physical sciences because many natural processes exhibit multiplicative variability and are strictly positive [9]. Limpert et al. [9] explain that when variability arises from the product of many independent factors such as cell division, metabolic rates, or tumour growth, the resulting distribution tends toward log-normal rather than normal. This makes log-normal sampling particularly suitable for parameters like growth rates and initial tumour sizes, which cannot be negative and are often right skewed.

### 2 Reinforcement learning implementation

We adapt the basic network architecture and hyperparameters used by Gallagher et al [6] and to work for NSCLC treated with erlotinib.

Coarse hyperparameter sweeps over learning rate, entropy coefficient, decay schedule, and ODE integration step were used to identify configurations that performed robustly across decision intervals after applying per-day scaling. Our unscaled values set at $d = 1$ were a learning rate $6 * 10^{-4}$, an entropy coefficient $10^{-3}$, a discount factor 0.99, gradient-norm clipping of 40, and an ODE step of 0.5 days. We used a learning rate decay value of 0.9 every 5000 steps. We also found that performance could be improved by scaling the reward, learning rate, gamma, and entropy coefficient according to the decision interval.

#### 2.1 Scaling across decision intervals

To ensure comparable behaviour across different decision intervals $d$, the following scaling was applied:

$$\begin{aligned} \gamma_{\text{step}} &= \gamma_{\text{day}}^{d}, \\ \beta_{\text{step}} &= \beta\, d, \\ \text{Learning\_rate}_{\text{step}} &= \frac{6 \times 10^{-4}}{\sqrt{d}}, \\ \text{Reward}_{\text{interval}} &= R_{\text{day}} \times d. \end{aligned}$$

These adjustments were chosen because they improved training stability and made policies more comparable across decision intervals. Entropy scaling was used to maintain exploration at longer decision intervals, where each action affects a longer simulated time window. The scaling was intended to reduce decision-interval dependence in the effective reward and optimisation dynamics.

#### 2.2 Network architecture

The network consisted of:

1. a 14-unit LSTM,
2. a dense feature extractor (128 $\rightarrow$ 64 $\rightarrow$ 32 $\rightarrow$ 16 $\rightarrow$ 10, ReLU),
3. a policy head (Softmax) producing action probabilities,
4. a value head (Linear) producing $V(s)$.

Policy weights were initialised to zero, giving a uniform initial policy. Global-norm gradient clipping at 40 was used to stabilise optimisation.

##### 2.3 Loss function

The loss function was:

$$\mathcal{L} = \frac{1}{4}\Big[\, \tfrac{1}{2}(V - R)^2 - \log \pi(a \mid s)\, \hat{A} - \beta\, H(\pi) \Big],$$

where the critic term fits $V(s)$ to observed returns, the policy-gradient term improves the policy in the direction of positive advantage $\hat{A}$, and the entropy term $H(\pi)$, adjusted by the entropy coefficient $\beta$, encourages exploration. The overall factor $\frac{1}{4}$ was retained from the original implementation and is implicitly accounted for in the tuned learning rate.

**Temporal information and model variants** The default policy input is only the current tumour burden ($w = 1$), and does not include information about past states. We tested augmenting the input with a longer history window of past decision-point observations (up to $w = 7$ previous $b(t)$ values) and with a simple trend feature (change in burden between decision points), but these did not yield consistent performance gains. We also tested simplified network variants (smaller feed-forward networks and removing the LSTM), but we ultimately retained the original setup including the LSTM feature extractor as even when using a single window without explicit temporal input, the recurrent hidden state can carry information forward across decision times within an episode, and in early tests we found this improved optimisation stability (details in Supplement 2). An episode is terminated at progression, or when the simulation horizon is reached. The maximum simulation horizon was set to 3528 days. Before between-interval analysis was performed all outcomes were cropped to a maximum of 3528 as this is a common multiple of all the decision intervals and allows fair comparisons to be made.

##### 2.4 Training and evaluation

**Training and evaluation procedure.** For the individual-policy experiments, a separate DRL agent was trained for each virtual patient and decision interval. Each policy was trained for 60,000 episodes across decision intervals $d \in \{1, 14, 28, 42, 56\}$ days. This range included shorter, idealised intervals to explore potential as well as longer $6 - 8$ week intervals typical of CT-based assessment in NSCLC. Training was performed using an Adam optimiser with an initial learning rate of $6 \times 10^{-4}$, an entropy coefficient $\beta = 10^{-3}$, and a discount factor $\gamma = 0.99$. The daily discount factor of 0.99 corresponds to an effective horizon of approximately 100 days. We retained this value, following Gallagher et al., because NSCLC dynamics are substantially faster than prostate cancer dynamics and because long-term outcomes are additionally encouraged through the long-term survival bonus. We also used exponential learning rate decay (factor 0.9 every 5000 steps). To ensure consistency across decision intervals, the learning rate, entropy, discounting factor, and reward magnitudes were scaled according to the decision interval $d$ (see Supplement 2.1 for details). The tumour dynamics were integrated with step size $\Delta t = 0.5$ days. Each patient-interval pair was trained using three independent random seeds, and the policy with the highest greedy-evaluation reward was retained.

After training, policies were evaluated using 300 independent stochastic rollouts per patient. For each patient–interval pair, we recorded time-to-progression (TTP) and the fraction of time under treatment. To obtain interpretable treatment rules, and to provide a representative final policy for protocol comparisons with deterministic policies, we also evaluated the greedy policy $a_t = \arg\max_a \pi_\theta(a \mid s_t)$. This greedy policy corresponds to the deterministic limit of the learned stochastic policy as exploration vanishes and thus reflects the effective outcome of training.

To reduce sensitivity to optimisation stochasticity and mitigate convergence to suboptimal local minima, each patient-interval pair was trained with three independent random seeds. Among these, we selected the policy where the greedy evaluation achieved the highest reward.

**Reward components and default values.** Values are reported on the per-day scale used in our implementation. These are based on the rewards used by Gallagher et al. with adjustments for NSCLC.

`survival_reward` **(0.1)** Reward for each day without progression.

`progression_penalty` **(-0.1)** Terminal penalty at progression.

`progression_multiplier` $(1.2\, x_{\text{total}}(0))$ Progression threshold.

`holiday_bonus` **(0.05)** Bonus for treatment holidays below threshold.

`holiday_penalty` **(-0.1)** Penalty for holidays above threshold.

`holiday_threshold` $(x_{\text{total}}(0))$ Safety threshold.

`bonus` **(5)** Long-term survival bonus.

`bonus_start_days` **(3 years)** Start time for long-term bonus.

#### 2.5 Policy interpretation via Monte Carlo boundary estimation

Because the learned policy is represented by a neural network, it does not provide an explicit treatment rule. Similar to Gallagher et al. [6], we make the decision rule interpretable by estimating an empirical treatment-propensity curve using Monte Carlo samples from stochastic evaluation rollouts. At each decision time we record the normalised tumour burden $b_t$ and the policy output $p_t = \pi_\theta(\mathrm{ON} \mid b_t)$. Pooling $(b_t, p_t)$ pairs across repeated rollouts provides a non-parametric estimate of treatment propensity as a function of tumour burden. To summarise this relationship, we partition the observed burden range into $n_{\mathrm{bins}}$ equal-width bins and compute the mean propensity within each bin. Uncertainty is visualised using within-bin percentile bands showing the 10th-90th percentiles of $p_t$. We can use this visualisation to extract the policy decision threshold, the tumour burden when the $p_t = 0.5$. The DRL policy can therefore be interpreted as "turn medication on only when the total tumour burden exceeds this threshold".

#### 2.6 Reward-shaping sweep

For the QoL-aware reward-shaping analysis, we performed an exploratory grid sweep over selected reward parameters in two representative virtual patients. The sweep was performed for decision intervals $d \in \{14, 28, 42\}$ days. For each patient, decision interval, and reward configuration, a separate DRL agent was trained for 50,000 episodes.

The reward parameters varied in the sweep were:

$$\texttt{holiday\_threshold} \in \{0.7, 0.8, 0.9, 1.0, 1.1, 1.2\},$$

$$\texttt{holiday\_penalty} \in \{-0.5, -0.1\},$$

$$\texttt{holiday\_bonus} \in \{0.05, 0.1\},$$

$$\texttt{survival\_reward} \in \{0.05, 0.1, 0.2\}.$$

The progression penalty was fixed at $\texttt{progression\_penalty} = -0.1$, and all other settings were held at their default values unless stated otherwise. This yielded $6 \times 2 \times 2 \times 3 = 72$ reward configurations per decision interval and patient, corresponding to 216 configurations per patient and 432 configurations across the two representative patients.

For each trained policy, we computed TTP, QAS, and mean QoL score $\mathrm{QAS}/T_{\mathrm{prog}}$. These results were used to explore trade-offs between survival, quality of life, and robustness across reward configurations.

Table S2: Tunable hyperparameters used for A2C training. Default values refer to the settings used in the main experiments unless otherwise stated.

| **Hyperparameter** | **Default** | **Description** |
|---|---|---|
| `learning_rate` | $6 \times 10^{-4}$ | Initial learning rate for the Adam optimiser (with exponential decay). |
| `decay_steps` | 5000 | Number of optimizer steps between learning rate decay events. |
| `decay_rate` | 0.9 | Multiplicative factor applied at each decay step (staircase schedule). |
| `gamma` | 0.99 | Discount factor for future rewards. |
| `entropy_coef` | 0.001 | Coefficient for entropy regularisation, encouraging exploration. |
| `entropy_anneal` | `False` | Whether to linearly anneal entropy from initial to final value. |
| `entropy_final` | 1e-5 | Final entropy coefficient (used only when annealing is enabled). |
| `entropy_anneal_frac` | 0.75 | Fraction of episodes over which entropy annealing occurs. |
| `window_size` | 1 | Number of past observations supplied to the LSTM (input window length). |
| `num_episodes` | 60000 | Number of training episodes. |
| `decision_interval` | $d \in \{1, 14, 28, 42, 56\}$ days | Time between dosing decisions made by the agent. |
| `target_days` | 3650 | Maximum simulation horizon used during training. Outcomes were cropped to 3528 days for analysis and comparison, as 3528 is a common multiple of all decision intervals. |
| `integration_step` | 0.5 days | Substep used for ODE integration within each decision interval. |
| `grad_clip_norm` | 40.0 | Global norm threshold for gradient clipping. |
| `restore_from` | `None` | Optional checkpoint for resuming training. |
| `output_dir` | `None` | Optional custom directory for saving results. |

### 3 Toxicity model

**Toxicity** ($T_n$)**.** We model treatment toxicity by tracking a normalised drug concentration $C_n$. In this scaling a full daily erlotinib dose of 150 mg increases $C_n$ by one unit. $C_n$ is not bounded above by 1 as repeated dosing can lead to drug accumulation. under continuous daily dosing with $t_{1/2} = 1.5$ days, the post-dose steady-state peak is approximately $1/(1 - e^{-0.462}) \approx 2.7$ units, and the corresponding pre-dose trough is approximately $2.7e^{-0.462} \approx 1.7$ units. Here $n$ indexes the internal simulation time-steps used to update the PK model, with $t_{n+1} = t_n + \Delta t$ and $C_n \approx C(t_n)$. $C_n$ is updated at every simulation time-step of length $\Delta t$ days. Erlotinib has an elimination half-life of around 36 hours (1.5 days) [4, 10]. We use a one-compartment model with first-order elimination in which absorption is approximated as instantaneous (each dose is treated as an impulse input), which has been shown to adequately describe erlotinib pharmacokinetics in NSCLC [5].

$$k_e = \frac{\ln 2}{t_{1/2}} = \frac{\ln 2}{1.5} \approx 0.462 \text{ day}^{-1}. \tag{1}$$

**Exact PK update used in the simulator.** Between dosing events, the PK model satisfies $\frac{dC}{dt} = -k_e C$, which has the closed-form solution

$$C(t) = C(t_{\text{last}})\, e^{-k_e\,(t - t_{\text{last}})},$$

where $t_{\text{last}}$ is the time of the most recent dose. In the simulator we use this exponential form directly between dose times (rather than numerically integrating the ODE) to avoid accumulating discretisation errors.

The treatment action $a$ is chosen once per decision interval $j$ of length $d$ days and is held fixed throughout that interval. If $a_j = 1$, doses are scheduled once per day; if $a_j = 0$, no doses are given. At a scheduled dosing time (when $a_j = 1$), we treat dosing as an instantaneous jump: we first evaluate the decayed concentration just before dosing using the exponential expression above, then add one unit (due to the normalisation), and reset $t_{\text{last}}$ to the new dosing time. We evaluate $C(t)$ on the internal time grid $t_n = n\Delta t$ and apply medication when a grid time-point matches a scheduled dose time (within the $\Delta t$ resolution).

**Mapping concentration to toxicity.** The dose-normalised concentration $C_n$ was mapped to a bounded toxicity burden using a standard sigmoidal $E_{\max}$ (Hill type) function with exponent $h = 2$. We set $E_{\max} = 1$ so that the resulting toxicity burden lies in $[0, 1]$ and can be used directly as the toxicity component in the QoL proxy:

$$B_{\text{tox}}(t_n) = \frac{C_n^2}{EC_{50}^2 + C_n^2}. \tag{2}$$

Here, $EC_{50}$ is the concentration at which $B_{\text{tox}}(t_n) = 0.5$. In all analyses reported here, we used $EC_{50} = 1$. The simulator allows $EC_{50}$ to vary across patients to represent inter-individual differences in toxicity sensitivity, but patient-specific toxicity sensitivity was not varied in the present study. Because treatment is binary in this model, toxicity closely follows whether medication is ON or OFF. We therefore treat toxicity as a simplified proxy for treatment-related side effects. We retain it as a separate QoL component because in future DRL settings with variable dosing or multiple drugs, toxicity and medication burden may no longer be equivalent.

**Defaults used in our simulator.** Unless stated otherwise, we use $\Delta t = 0.1$ days, $t_{1/2} = 1.5$ days, $EC_{50} = 1.0$, $h = 2$, and initialise $C_0 = 0$. Dosing is scheduled once per day when ON and no dose is applied when OFF. For each decision interval, we use the end-of-interval toxicity value as the toxicity burden component.

### 4 Time to progression summary statistics

Table S3: Summary statistics for time to progression (TTP, months) by treatment policy and decision interval.

| Policy | Decision (days) | Mean | SD | Median | IQR |
|---|---|---|---|---|---|
| MTD | 1 | 14.73 | 5.17 | 15.11 | (10.95, 18.53) |
| | 14 | 14.73 | 5.17 | 15.11 | (10.95, 18.53) |
| | 28 | 14.73 | 5.17 | 15.11 | (10.95, 18.53) |
| | 42 | 14.73 | 5.17 | 15.11 | (10.95, 18.53) |
| | 56 | 14.73 | 5.17 | 15.11 | (10.95, 18.53) |
| DRL agent | 1 | 61.83 | 43.76 | 49.59 | (19.11, 115.78) |
| | 14 | 40.14 | 28.53 | 35.66 | (18.86, 52.59) |
| | 28 | 29.00 | 18.99 | 25.85 | (15.48, 37.85) |
| | 42 | 23.61 | 13.24 | 21.42 | (13.02, 30.59) |
| | 56 | 20.22 | 9.93 | 18.95 | (11.54, 26.95) |
| Zhang et al. protocol | 1 | 49.21 | 37.12 | 40.41 | (21.99, 65.09) |
| | 14 | 36.87 | 31.92 | 29.46 | (11.43, 53.58) |
| | 28 | 22.45 | 27.22 | 7.31 | (3.31, 35.86) |
| | 42 | 17.48 | 20.36 | 5.75 | (3.81, 28.76) |
| | 56 | 12.67 | 13.23 | 5.79 | (4.38, 17.53) |

### 5 Kaplan Meier plots with risk tables for all decision intervals

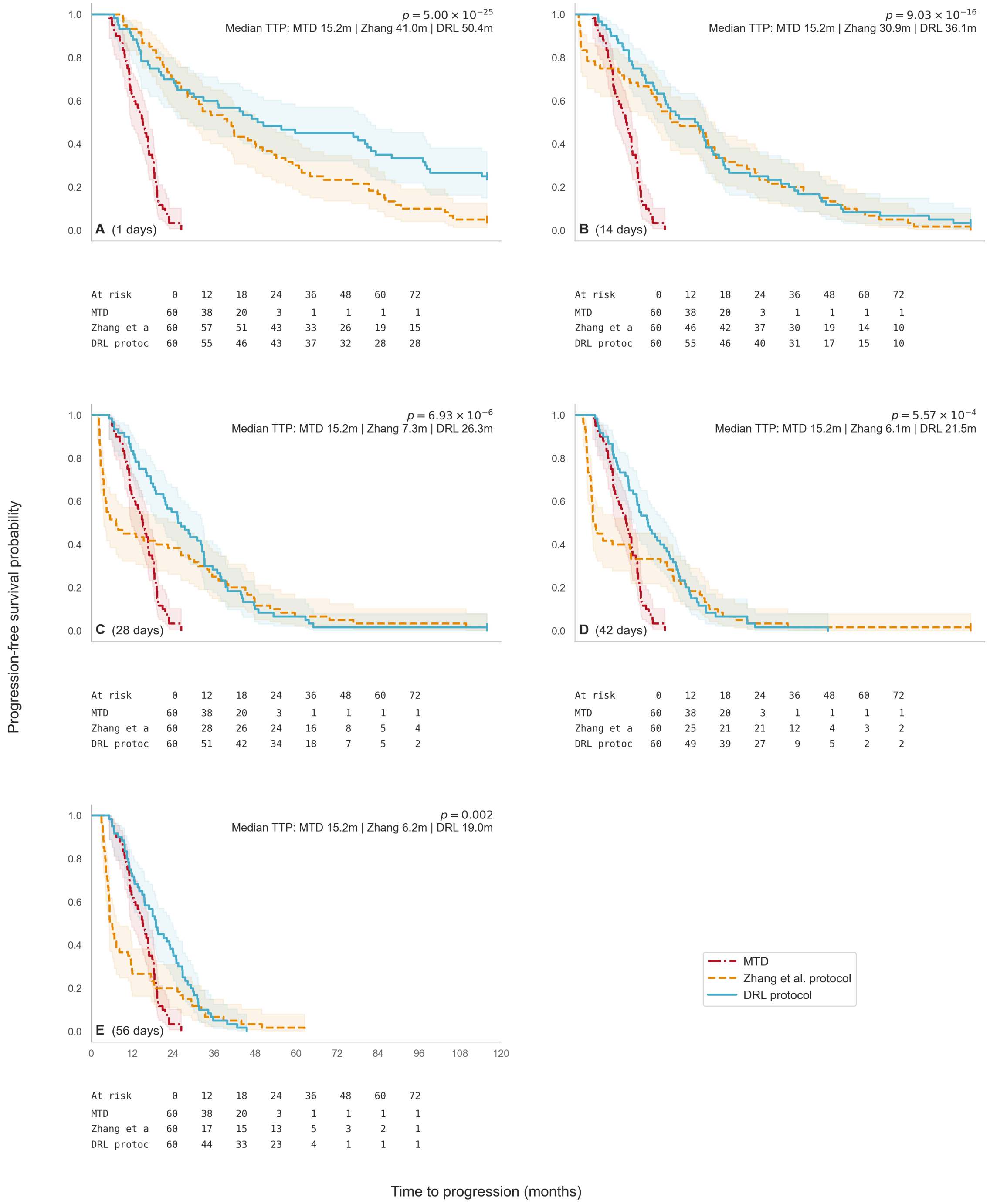


Figure S2: Kaplan Meier plots showing probability of progression over time for the MTD, Zhang et al., and DRL protocols. Risk tables for each subplot indicate the number not yet progressed (still at risk) at each time.

#### 5.1 Pairwise log-rank tests by decision interval

Table S4: Pairwise log-rank tests by decision interval. Reported values are adjusted $p$-values with significance coding.

(a) Bonferroni adjustment (within each decision interval; three pairwise tests per panel).

| Decision interval (days) | MTD vs Zhang et al. | MTD vs DRL | Zhang et al. vs DRL |
|---|---|---|---|
| 1 | $1.76 \times 10^{-18}$ **** | $2.97 \times 10^{-15}$ **** | $4.21 \times 10^{-2}$ * |
| 14 | $9.02 \times 10^{-10}$ **** | $6.34 \times 10^{-15}$ **** | $1.00 \times 10^{0}$ ns |
| 28 | $1.01 \times 10^{-1}$ ns | $1.42 \times 10^{-10}$ **** | $6.92 \times 10^{-1}$ ns |
| 42 | $7.12 \times 10^{-1}$ ns | $1.58 \times 10^{-7}$ **** | $3.94 \times 10^{-1}$ ns |
| 56 | $1.00 \times 10^{0}$ ns | $2.49 \times 10^{-5}$ **** | $2.45 \times 10^{-2}$ * |

(b) Holm–Bonferroni adjustment (within each decision interval; three pairwise tests per panel). Holm–Bonferroni produced the same significance categories as Bonferroni under the reporting thresholds.

| Decision interval (days) | MTD vs Zhang | MTD vs DRL | Zhang vs DRL |
|---|---|---|---|
| 1 | $1.76 \times 10^{-18}$ **** | $1.98 \times 10^{-15}$ **** | $1.40 \times 10^{-2}$ * |
| 14 | $6.01 \times 10^{-10}$ **** | $6.34 \times 10^{-15}$ **** | $6.21 \times 10^{-1}$ ns |
| 28 | $6.76 \times 10^{-2}$ ns | $1.42 \times 10^{-10}$ **** | $2.31 \times 10^{-1}$ ns |
| 42 | $2.62 \times 10^{-1}$ ns | $1.58 \times 10^{-7}$ **** | $2.62 \times 10^{-1}$ ns |
| 56 | $4.25 \times 10^{-1}$ ns | $2.49 \times 10^{-5}$ **** | $1.64 \times 10^{-2}$ * |

Significance coding: ns ($p \geq 0.05$), * ($p < 0.05$), ** ($p < 0.01$), *** ($p < 0.001$), **** ($p < 10^{-4}$).

### 6 Medication on percentage table

Table S5: Mean and median percentages of time that medication is on under the DRL policy for 60 virtual patients by decision interval (days). Medication usage is calculated over the treatment period, either until progression or when the time horizon of 3528 days is reached

| | 1 | 14 | 28 | 42 | 56 |
|---|---|---|---|---|---|
| Mean | 30.0 | 35.9 | 48.0 | 57.6 | 65.2 |
| Median | 19.9 | 32.2 | 42.7 | 53.8 | 59.1 |

### 7 Quality adjusted survival

#### 7.1 QAS Summary tables

Table S6: Median and inter-quartile range (IQR) for time-to-progression (TTP), quality-adjusted survival (QAS), and QAS/day by preference profile, decision interval, and protocol

| Profile | $d$ (days) | Protocol | TTP [IQR] | QAS [IQR] | QAS/day [IQR] |
|---|---|---|---|---|---|
| Balanced | 1 | MTD | 460 [333–564] | 142 [98–177] | 0.310 [0.303–0.319] |
| | 1 | DRL Agent | 1509 [582–3524] | 914 [333–2184] | 0.609 [0.543–0.643] |
| | 1 | Zhang Protocol | 1230 [669–1981] | 709 [390–1235] | 0.600 [0.571–0.655] |
| | 14 | MTD | 460 [333–564] | 142 [98–177] | 0.310 [0.303–0.319] |
| | 14 | DRL Agent | 1085 [574–1601] | 657 [332–1030] | 0.615 [0.566–0.674] |
| | 14 | Zhang Protocol | 897 [348–1631] | 550 [175–1042] | 0.641 [0.591–0.666] |
| | 28 | MTD | 460 [333–564] | 142 [98–177] | 0.310 [0.303–0.319] |
| | 28 | DRL Agent | 787 [471–1152] | 487 [250–709] | 0.597 [0.527–0.647] |
| | 28 | Zhang Protocol | 223 [101–1092] | 153 [65–683] | 0.653 [0.601–0.690] |
| | 42 | MTD | 460 [333–564] | 142 [98–177] | 0.310 [0.303–0.319] |
| | 42 | DRL Agent | 652 [396–931] | 350 [202–549] | 0.556 [0.499–0.620] |
| | 42 | Zhang Protocol | 175 [116–876] | 116 [76–605] | 0.655 [0.603–0.707] |
| | 56 | MTD | 460 [333–564] | 142 [98–177] | 0.310 [0.303–0.319] |
| | 56 | DRL Agent | 577 [351–820] | 303 [154–508] | 0.536 [0.411–0.562] |
| | 56 | Zhang Protocol | 176 [133–534] | 120 [85–336] | 0.651 [0.590–0.702] |
| Toxicity-adverse | 1 | MTD | 460 [333–564] | 100 [71–121] | 0.218 [0.214–0.220] |
| | 1 | DRL Agent | 1509 [582–3524] | 1161 [327–2784] | 0.757 [0.620–0.816] |
| | 1 | Zhang Protocol | 1230 [669–1981] | 815 [450–1440] | 0.690 [0.657–0.761] |
| | 14 | MTD | 460 [333–564] | 100 [71–121] | 0.218 [0.214–0.220] |
| | 14 | DRL Agent | 1085 [574–1601] | 712 [357–1144] | 0.689 [0.593–0.747] |
| | 14 | Zhang Protocol | 897 [348–1631] | 624 [196–1188] | 0.729 [0.678–0.759] |
| | 28 | MTD | 460 [333–564] | 100 [71–121] | 0.218 [0.214–0.220] |
| | 28 | DRL Agent | 787 [471–1152] | 502 [252–772] | 0.637 [0.522–0.691] |
| | 28 | Zhang Protocol | 223 [101–1092] | 168 [71–770] | 0.736 [0.664–0.758] |
| | 42 | MTD | 460 [333–564] | 100 [71–121] | 0.218 [0.214–0.220] |
| | 42 | DRL Agent | 652 [396–931] | 335 [198–573] | 0.552 [0.480–0.636] |
| | 42 | Zhang Protocol | 175 [116–876] | 123 [79–649] | 0.702 [0.643–0.748] |
| | 56 | MTD | 460 [333–564] | 100 [71–121] | 0.218 [0.214–0.220] |
| | 56 | DRL Agent | 577 [351–820] | 287 [123–495] | 0.517 [0.335–0.562] |
| | 56 | Zhang Protocol | 176 [133–534] | 122 [86–347] | 0.693 [0.617–0.731] |
| Visit-adverse | 1 | MTD | 460 [333–564] | 8 [6–10] | 0.018 [0.018–0.018] |
| | 1 | DRL Agent | 1509 [582–3524] | 251 [63–595] | 0.163 [0.124–0.175] |
| | 1 | Zhang Protocol | 1230 [669–1981] | 167 [94–302] | 0.142 [0.134–0.158] |
| | 14 | MTD | 460 [333–564] | 350 [253–429] | 0.760 [0.760–0.761] |
| | 14 | DRL Agent | 1085 [574–1601] | 956 [491–1426] | 0.883 [0.860–0.895] |
| | 14 | Zhang Protocol | 897 [348–1631] | 792 [297–1442] | 0.892 [0.879–0.900] |
| | 28 | MTD | 460 [333–564] | 363 [263–445] | 0.789 [0.789–0.789] |
| | 28 | DRL Agent | 787 [471–1152] | 707 [408–1038] | 0.893 [0.865–0.908] |
| | 28 | Zhang Protocol | 223 [101–1092] | 206 [93–991] | 0.919 [0.903–0.925] |
| | 42 | MTD | 460 [333–564] | 367 [266–450] | 0.799 [0.799–0.799] |
| | 42 | DRL Agent | 652 [396–931] | 578 [349–851] | 0.882 [0.865–0.902] |
| | 42 | Zhang Protocol | 175 [116–876] | 160 [106–820] | 0.920 [0.906–0.929] |
| | 56 | MTD | 460 [333–564] | 369 [268–453] | 0.803 [0.803–0.803] |
| | 56 | DRL Agent | 577 [351–820] | 494 [301–748] | 0.877 [0.833–0.890] |
| | 56 | Zhang Protocol | 176 [133–534] | 164 [121–495] | 0.922 [0.902–0.929] |
| Tumour-burden-adverse | 1 | MTD | 460 [333–564] | 283 [198–354] | 0.619 [0.602–0.640] |
| | 1 | DRL Agent | 1509 [582–3524] | 493 [290–899] | 0.353 [0.295–0.530] |
| | 1 | Zhang Protocol | 1230 [669–1981] | 541 [294–899] | 0.449 [0.430–0.461] |
| | 14 | MTD | 460 [333–564] | 283 [198–354] | 0.619 [0.602–0.640] |
| | 14 | DRL Agent | 1085 [574–1601] | 538 [306–829] | 0.508 [0.476–0.580] |
| | 14 | Zhang Protocol | 897 [348–1631] | 427 [150–801] | 0.482 [0.456–0.516] |
| | 28 | MTD | 460 [333–564] | 283 [198–354] | 0.619 [0.602–0.640] |
| | 28 | DRL Agent | 787 [471–1152] | 485 [273–633] | 0.594 [0.564–0.636] |
| | 28 | Zhang Protocol | 223 [101–1092] | 129 [54–591] | 0.549 [0.506–0.598] |
| | 42 | MTD | 460 [333–564] | 283 [198–354] | 0.619 [0.602–0.640] |
| | 42 | DRL Agent | 652 [396–931] | 439 [260–578] | 0.626 [0.589–0.666] |
| | 42 | Zhang Protocol | 175 [116–876] | 110 [76–529] | 0.597 [0.555–0.650] |
| | 56 | MTD | 460 [333–564] | 283 [198–354] | 0.619 [0.602–0.640] |
| | 56 | DRL Agent | 577 [351–820] | 369 [237–556] | 0.649 [0.615–0.679] |
| | 56 | Zhang Protocol | 176 [133–534] | 117 [87–342] | 0.633 [0.591–0.682] |

#### 7.2 QAS Deltas (Differences between profile pairs) across decision intervals

Table S7: Median paired ΔQAS [25th, 75th percentiles] by protocol comparison, decision interval, and preference profile. Positive values indicate greater QAS under the first-named protocol. QAS differences are reported in days.

| Comparison | Decision (days) | Preference profile | | | |
|---|---|---|---|---|---|
| | | Balanced | Toxicity-adverse | Tumour-burden-adverse | Visit-adverse |
| DRL – MTD | 1 | 740 [211, 1999] | 1041 [246, 2643] | 157 [75, 571] | 241 [57, 583] |
| | 14 | 527 [233, 866] | 615 [294, 1035] | 205 [97, 487] | 570 [236, 1031] |
| | 28 | 309 [160, 520] | 377 [188, 643] | 171 [75, 344] | 344 [157, 632] |
| | 42 | 213 [100, 369] | 249 [127, 445] | 136 [54, 248] | 215 [84, 372] |
| | 56 | 172 [34, 323] | 199 [43, 374] | 88 [7, 186] | 142 [5, 302] |
| Zhang – MTD | 1 | 557 [291, 1076] | 718 [382, 1330] | 182 [79, 559] | 160 [88, 293] |
| | 14 | 415 [88, 874] | 529 [132, 1069] | 153 [−16, 458] | 460 [79, 993] |
| | 28 | 28 [−43, 520] | 72 [−1, 653] | −53 [−160, 252] | −54 [−195, 585] |
| | 42 | 15 [−32, 434] | 50 [0, 528] | −61 [−151, 208] | −68 [−179, 387] |
| | 56 | 0 [−49, 216] | 29 [−1, 265] | −75 [−199, 78] | −87 [−256, 188] |
| DRL – Zhang | 1 | 84 [−55, 598] | 156 [−86, 858] | 11 [−51, 36] | 50 [−19, 192] |
| | 14 | 5 [−58, 210] | −9 [−88, 224] | 24 [2, 117] | 4 [−86, 313] |
| | 28 | 32 [−70, 263] | 15 [−97, 279] | 74 [−1, 269] | 88 [−90, 385] |
| | 42 | 45 [−90, 215] | 28 [−127, 205] | 109 [−12, 269] | 136 [−69, 362] |
| | 56 | 45 [−66, 275] | 21 [−86, 252] | 98 [−1, 330] | 132 [−38, 449] |

Table S8: P-values from exact two-sided sign tests on paired patient-level QAS differences by protocol comparison, decision interval, and preference profile. For each comparison, the test was applied to the non-zero within-patient QAS differences between the two named protocols, using a binomial test with null probability 0.5. Superscripts indicate significance levels: $^{*}p < 0.05$, $^{**}p < 0.01$.

| Comparison | Decision (days) | Preference profile | | | |
|---|---|---|---|---|---|
| | | Balanced | Toxicity-adverse | Tumour-burden-adverse | Visit-adverse |
| DRL – MTD | 1 | $<0.001^{**}$ | $<0.001^{**}$ | $<0.001^{**}$ | $<0.001^{**}$ |
| | 14 | $<0.001^{**}$ | $<0.001^{**}$ | $<0.001^{**}$ | $<0.001^{**}$ |
| | 28 | $<0.001^{**}$ | $<0.001^{**}$ | $<0.001^{**}$ | $<0.001^{**}$ |
| | 42 | $<0.001^{**}$ | $<0.001^{**}$ | $<0.001^{**}$ | $<0.001^{**}$ |
| | 56 | $<0.001^{**}$ | $<0.001^{**}$ | $<0.001^{**}$ | $<0.001^{**}$ |
| Zhang – MTD | 1 | $<0.001^{**}$ | $<0.001^{**}$ | $<0.001^{**}$ | $<0.001^{**}$ |
| | 14 | $<0.001^{**}$ | $0.001^{**}$ | $<0.001^{**}$ | $<0.001^{**}$ |
| | 28 | 0.897 | 0.519 | 0.519 | 0.519 |
| | 42 | 0.245 | 0.519 | 0.519 | 0.699 |
| | 56 | 1.000 | $0.013^{*}$ | $0.027^{*}$ | $0.027^{*}$ |
| DRL – Zhang | 1 | 0.155 | 0.052 | 0.155 | 0.052 |
| | 14 | 0.897 | $<0.001^{**}$ | 0.897 | 0.897 |
| | 28 | 0.245 | $<0.001^{**}$ | 0.519 | 0.519 |
| | 42 | 0.366 | $0.006^{**}$ | 0.067 | 0.067 |
| | 56 | 0.092 | $<0.001^{**}$ | $0.002^{**}$ | $0.002^{**}$ |

Table S9: Patient-level QAS win rates (%, $n = 60$) by protocol comparison, decision interval, and preference profile. Values indicate the percentage of patients where the first-named protocol achieved higher QAS than the second.

| Comparison | Decision (days) | Preference profile | | | |
|---|---|---|---|---|---|
| | | Balanced | Toxicity-adverse | Tumour-burden-adverse | Visit-adverse |
| DRL > MTD | 1 | 100.0 | 100.0 | 98.3 | 98.3 |
| | 14 | 100.0 | 100.0 | 98.3 | 100.0 |
| | 28 | 96.7 | 96.7 | 96.7 | 95.0 |
| | 42 | 91.7 | 91.7 | 91.7 | 90.0 |
| | 56 | 83.3 | 83.3 | 83.3 | 80.0 |
| Zhang > MTD | 1 | 100.0 | 100.0 | 95.0 | 100.0 |
| | 14 | 78.3 | 80.0 | 71.7 | 76.7 |
| | 28 | 51.7 | 75.0 | 45.0 | 45.0 |
| | 42 | 58.3 | 73.3 | 45.0 | 46.7 |
| | 56 | 50.0 | 73.3 | 33.3 | 35.0 |
| DRL > Zhang | 1 | 60.0 | 63.3 | 60.0 | 63.3 |
| | 14 | 51.7 | 48.3 | 76.7 | 51.7 |
| | 28 | 58.3 | 55.0 | 75.0 | 55.0 |
| | 42 | 56.7 | 55.0 | 68.3 | 61.7 |
| | 56 | 61.7 | 58.3 | 73.3 | 70.0 |

### References

[1] Renée Brady and Heiko Enderling. 2019. Mathematical Models of Cancer: When to Predict Novel Therapies, and When Not to. Bulletin of Mathematical Biology 81, 9 (2019), 3722–3731. doi:10.1007/s11538-019-00640-x

[2] Jessica J. Cunningham, Frank Thuijsman, Ralf Peeters, Yannick Viossat, Joel S. Brown, Robert A. Gatenby, and Kateřina Staňková. 2020. Optimal control to reach eco-evolutionary stability in metastatic castrate-resistant prostate cancer. PLOS ONE 15, 12 (Dec. 2020), e0243386. doi:10.1371/journal.pone.0243386

[3] Marisa C. Eisenberg and Harsh V. Jain. 2017. A confidence building exercise in data and identifiability: Modeling cancer chemotherapy as a case study. Journal of Theoretical Biology 431 (2017), 63–78. doi:10.1016/j.jtbi.2017.07.018

[4] European Medicines Agency. 2023. Tarceva (erlotinib) EPAR—Product Information (Summary of Product Characteristics). https://www.ema.europa.eu/en/documents/product-information/tarceva-epar-product-information_en.pdf. Section 5.2 Pharmacokinetic properties; revised May 2023.

[5] Evelina Cardoso, Monia Guidi, Nihel Khoudour, Pascaline Boudou-Rouquette, Elizabeth Fabre, Camille Tlemsani, Jennifer Arrondeau, François Goldwasser, Michel Vidal, Marie Paule Schneider, Anna Dorothea Wagner, Nicolas Widmer, Benoit Blanchet, and Chantal Csajka. 2020. Population Pharmacokinetics of Erlotinib in Patients With Non–small Cell Lung Cancer: Its Application for Individualized Dosing Regimens in Older Patients. Clinical Therapeutics 42, 7 (2020), 1302–1316. doi:10.1016/j.clinthera.2020.05.008

[6] Kit Gallagher, Maximilian A.R. Strobl, Derek S. Park, Fabian C. Spoendlin, Robert A. Gatenby, Philip K. Maini, and Alexander R.A. Anderson. 2024. Mathematical Model-Driven Deep Learning Enables Personalized Adaptive Therapy. Cancer Research 84, 11 (2024), 1929–1941. doi:10.1158/0008-5472.CAN-23-2040

[7] Simon P. Hart, Robert P. Freckleton, and Jonathan M. Levine. 2018. How to quantify competitive ability. Journal of Ecology 106, 5 (2018), 1902–1909. doi:10.1111/1365-2745.12954

[8] Laura R. Jansén-Storbacka, Kailas S. Honasoge, Eva Molnárová, Arina Soboleva, Bram C. Agema, Marthe S. Paats, Dirk Jan A. R. Moes, G. D. Marijn Veerman, Alethea B. T. Barbaro, Roel Dobbe, Irene Grossmann, Sepinoud Azimi, Ron H. J. Mathijssen, Anne-Marie C. Dingemans, and Kateřina

Staňková. 2026. Can evolutionary therapy be applied in non-small cell lung cancer? Scientific Reports 16, 1 (2026), 36712. doi:10.1038/s41598-026-36712-x

[9] Eckhard Limpert, Werner A. Stahel, and Markus Abbt. 2001. Log-normal Distributions across the Sciences: Keys and Clues. BioScience 51, 5 (2001), 341–352. doi:10.1641/0006-3568(2001)051[0341:LNDATS]2.0.CO;2

[10] J. F. Lu, S. M. Eppler, J. Wolf, M. Hamilton, A. Rakhit, R. Bruno, and B. L. Lum. 2006. Clinical pharmacokinetics of erlotinib in patients with solid tumors and exposure-safety relationship in patients with non-small cell lung cancer. Clinical Pharmacology & Therapeutics 80, 2 (2006), 136–145. doi:10.1016/j.clpt.2006.04.007

[11] J. Christopher D. Terry. 2024. Uncertain competition coefficients undermine inferences about coexistence. Nature 632, 8027 (2024), E9–E14. doi:10.1038/s41586-023-06859-y
